# TDP-43-M323K causes abnormal brain development and progressive cognitive and motor deficits associated with mislocalised and increased levels of TDP-43

**DOI:** 10.1101/2023.12.27.573393

**Authors:** Juan M. Godoy-Corchuelo, Zeinab Ali, Jose M. Brito Armas, Aurea B. Martins-Bach, Irene García-Toledo, Luis C. Fernández-Beltrán, Juan I. López-Carbonero, Pablo Bascuñana, Shoshana Spring, Irene Jimenez-Coca, Ramón A. Muñoz de Bustillo Alfaro, Maria J. Sánchez-Barrena, Remya R. Nair, Brian J. Nieman, Jason P. Lerch, Karla L. Miller, Hande P Ozdinler, Elizabeth M.C. Fisher, Thomas J. Cunningham, Abraham Acevedo-Arozena, Silvia Corrochano

**Affiliations:** Neurological Disorders Group, Hospital Clínico San Carlos, Instituto de Investigación Sanitaria Hospital Clínico San Carlos (IdiSSC), Madrid 28040, Spain; MRC Harwell Institute, Oxfordshire, UK; Unidad de Investigación, Hospital Universitario de Canarias, ITB-ULL and CIBERNED, La Laguna, Spain; Wellcome Centre for Integrative Neuroimaging, University of Oxford, Oxford, UK; Department of Medicine, Universidad Complutense de Madrid, Madrid, Spain; Brain Mapping Group, Hospital Clínico San Carlos, IdISSC, Madrid, Spain; Mouse Imaging Centre, The Hospital for Sick Children, Toronto, ON, Canada; Department of Crystallography and Structural Biology. Institute of Physical Chemistry “Blas Cabrera”, CSIC, Madrid, Spain; Nucleic Acid Therapy Accelerator (NATA), Harwell Campus, Oxfordshire, UK; Department of Neurology, Feinberg School of Medicine, Northwestern University, Chicago, IL, USA; Department of Neuromuscular Diseases, and UCL Queen Square Motor Neuron Disease Centre, UCL. Institute of Neurology, London, UK; MRC Prion Unit at UCL, UCL Institute of Prion Diseases, London, UK

**Author notes:** these authors contributed equally to this work (Juan M. Godoy-Corchuelo, Zeinab Ali, Jose M. Brito Armas). these authors contributed equally to this work (Thomas J. Cunningham, Abraham Acevedo-Arozena, Silvia Corrochano).

**Keywords:** TDP-43, Cognitive alterations, Motor disturbances, ALS-FTD, TDP-43 Proteinopathies, Development

## Abstract

TDP-43 pathology is found in several neurodegenerative disorders, collectively referred to as “TDP-43 proteinopathies”. Aggregates of TDP-43 are present in the brains and spinal cords of >97% of amyotrophic lateral sclerosis (ALS), and in brains of ∼50% of frontotemporal dementia (FTD) patients. While mutations in the TDP-43 gene (*TARDBP*) are usually associated with ALS, many clinical reports have linked these mutations to cognitive impairments and/or FTD, but also to other neurodegenerative disorders including Parkinsonism (PD) or progressive supranuclear palsy (PSP). TDP-43 is a ubiquitously expressed, highly conserved RNA-binding protein that is involved in many cellular processes, mainly RNA metabolism. To investigate systemic pathological mechanisms in TDP-43 proteinopathies, aiming to capture the pleiotropic effects of TDP-43 mutations, we have further characterised a mouse model carrying a point mutation (M323K) within the endogenous *Tardbp* gene. Homozygous mutant mice developed cognitive and behavioural deficits as early as 3 months of age. This was coupled with significant brain structural abnormalities, mainly in the cortex, hippocampus, and white matter fibres, together with progressive cortical interneuron degeneration and neuroinflammation. At the motor level, progressive phenotypes appeared around 6 months of age. Thus, cognitive phenotypes appeared to be of a developmental origin with a mild associated progressive neurodegeneration, while the motor and neuromuscular phenotypes seemed neurodegenerative, underlined by a progressive loss of upper and lower motor neurons as well as distal denervation. This is accompanied by progressive elevated TDP-43 protein and mRNA levels in cortex and spinal cord of homozygous mutant mice from 3 months of age, together with increased cytoplasmic TDP-43 mislocalisation in cortex, hippocampus, hypothalamus, and spinal cord at 12 months of age. In conclusion, we find that *Tardbp* M323K homozygous mutant mice model many aspects of human TDP-43 proteinopathies, evidencing a dual role for TDP-43 in brain morphogenesis as well as in the maintenance of the motor system, making them an ideal *in vivo* model system to study the complex biology of TDP-43.

## Introduction

The transactivation response element DNA-binding protein of 43 KDa (TDP-43) is a 414-residue heterogeneous ribonucleoprotein (hnRNP) that in normal conditions is highly enriched in the nucleus, although it can shuttle between the nucleus and the cytoplasm. The *TARDBP* gene encodes this highly conserved protein, which consists of two RNA recognition motifs and a C-terminal glycine-rich domain, with nuclear localisation and export signals that are critical for its nuclear-cytoplasmic shuttling. In the nucleus, TDP-43 is found in different nuclear bodies with an important function in modulating RNA processing, particularly alternative splicing through direct mRNA binding to hundreds of transcripts. The roles of TDP-43 in the cytoplasm are less clear; it can shuttle to the cytoplasm in response to different stresses, colocalizing with stress granules, although TDP-43 clearly has other physiological functions outside the nucleus, such as in myogenesis in which cytoplasmic TDP-43 granules are required for correct muscle differentiation (Vogler et al., 2018) and in the regulation of synapsis (S. Ling, 2018). However, in TDP-43 pathology, the normally nuclear TDP-43 protein is mislocalized to the cytoplasm where it can subsequently form poly-phosphorylated and polyubiquitinated insoluble inclusions, leading to the loss of nuclear functions such as in splicing regulation, as well as potential gain of cytoplasmic toxicity (Buratti & Baralle, 2008). Despite the recent advancement in the knowledge of the role of TDP-43 in RNA regulation in neurons and glia, there are still many unknowns about its normal roles in the broader neuronal system of healthy ageing and in the multiple associated diseases.

TDP-43 proteinopathies are a heterogeneous group of neurodegenerative diseases that share the pathological presence of mislocalization and aggregation of the protein TDP-43, among other proteins and molecules, in different cellular populations and regions of the central nervous system (CNS) (de Boer et al., 2021). Pathogenic deposits of TDP-43 have been identified in postmortem samples in different brain regions and spinal cord of an increasing number of neurodegenerative disorders, including the great majority of ALS cases, around 50% of all FTD cases, and all LATE cases (limbic-predominant age-related TDP-43 encephalopathy) (Nelson et al., 2019), a new dementia entity that overlaps with Alzheimer’s disease (AD) and occurs particularly in old age. Moreover, secondary TDP-43 pathology has been found in other disorders including Alzheimer’s disease (AD), Lewy body disease (LBD), argyrophilic grain disease (AGD), Parkinsonism or Huntington’s disease (Amador-Ortiz et al., 2007; Chanson et al., 2010; Davidson et al., 2009; Josephs et al., 2014; Kabashi et al., 2008; Schwab et al., 2008; Sreedharan et al., 2008). Interestingly, TDP-43 proteinopathy also occurs in the brain of healthy and cognitively normal aged individuals (Arnold et al., 2013; Uchino et al., 2015). Overall, the wide variety of TDP-43 proteinopathies shows that TDP-43 has a broad role in the CNS beyond the maintenance of the motor system, particularly in cognition (Kawakami et al., 2019). Thus, understanding the multiple roles of TDP-43 in the different brain and spinal cord areas, and in different stages of life, is essential to help explain the heterogeneous clinical outcomes and associated neurodegenerative diseases that present as TDP-43 proteinopathies.

Although the great majority of TDP-43 proteinopathies are sporadic cases, or caused by mutations in other genes (e.g., in the *C9Orf72* and *GRN* genes), some cases are caused by mutations in *TARDBP* itself. To date, most mutations have been found in ALS patients, although mutations haven also been described in FTD, ALS with FTD, as well as Parkinsonism or progressive supranuclear palsy (Cannas et al., 2013; Caroppo et al., 2016; Chen et al., 2021; Kovacs et al., 2009; Tiloca et al., 2022). The mechanisms by which TDP-43 mutations lead to the death of specific neuronal pools remains unknown. Almost all causative mutations in TDP-43 lie within its C-terminal glycine-rich region, where changes to the prion-like domain can affect its aggregation potential. Although the majority of mutations are dominant, there are also reports of homozygous *TARDBP* mutations leading to ALS, FTD and also other disorders (Borghero et al., 2011; Mosca et al., 2012). At the molecular level, we and others have shown that different C-terminal TDP-43 mutations, such as Q331K and M323K, can lead to a splicing gain of function when expressed at physiological levels (Fratta et al., 2018). However, in TDP-43 pathology, TDP-43 exits the nucleus, leading to the loss of nuclear function including in splicing regulation and the appearance of pathological events such as cryptic exons that seem to define at least the end-stage of TDP-43 proteinopathies (Cao et al., 2023; J. P. Ling et al., 2015; Ma et al., 2022).

TDP-43 pathology has been found in different cellular types and in different clinical outcomes and neurodegenerative disorders (Kawakami et al., 2019). As there are a variety of disorders with TDP-43 pathology encompassing different brain and spinal cord regions, this suggests that TDP-43 has a broader role in the CNS, beyond the maintenance of the motor system. Different patterns of TDP-43 pathology have been associated with different clinical entities. For example, TDP-43 deposits in ALS are in the spinal cord and areas of the frontal and temporal lobes. In FTD, TDP-43 is found more in the orbitofrontal cortex, also in the temporal and frontal cortex, visual and cerebellum, in addition to progressing to the motor area (Kawakami et al., 2019). In LATE, the pathology is located in the hippocampus and amygdala. Therefore, diverse TDP-43 pathology exists, variably affecting multiple areas of the brain (Jo et al., 2020), while the functions of TDP-43 across different cell types of the central and peripheral nervous system is poorly understood and requires further investigation.

RNA binding proteins such as TDP-43 are exquisitely dosage-sensitive, with an auto-regulatory pathway controlling their tight regulation through a mechanism involving TDP-43 protein binding to the 3’UTR of its own transcript (Avendaño-Vázquez et al., 2012). Thus, overexpression or downregulation of TDP-43 leads to toxicity in a variety of model systems. Therefore, to accurately study TDP-43 biology and its central role in the related proteinopathies, it is critical to work with models that maintain physiological control of gene expression. Over the past few years, we and others have been working with knock-in (KI) lines carrying point mutations within the mouse *Tardbp* gene, including the ENU-induced M323K mutation, which in homozygosity develops progressive motor neuron degeneration underlined with a splicing gain of function (Fratta et al., 2018). While this mutation has not been identified in human patients, it resides within the prion-like domain where various human ALS-causing mutations have been found.

Here, we further characterise the pleiotropic effects of the M323K point mutation longitudinally to study the implications of expressing the mutant protein physiologically, along the entire lifespan of the mouse and its effects in different systems. This offers the unique opportunity of evaluating the progressive nature of some of the observed phenotypes, uncovering novel functions for TDP-43 in brain morphogenesis and maintenance whilst illuminating its role in disease pathogenesis.

## Material and Methods

### Animals

#### Housing conditions and licensing

Mice were kept in auto-ventilated cages and fed *ad libitum* (Rat and Mouse Breeding 3 (RM3), Special Diet Services) with free access to water. Mice were kept under constant conditions with a regular 12 hour light and dark cycle, temperature of 21 ± 2 °C and humidity 55 ± 10 %. Mice were housed in same sex cages of up to five mice. Cages contained Aspen Chips bedding (Datesand), shredded paper and a rodent tunnel for enrichment. The animals were always kept in their cages except in the case of a behavioural test. Isolation time was kept as short as possible so as not to affect the results of other behavioural tests. Separated male and female cohorts were used in most of the analyses unless specified. All behavioural tests were conducted blind to the operator.

All mice were maintained and studied according to international, national and regional guidelines. At MRC Harwell, UK Home Office legislation was followed (Home Office Project Licence 20/0005), with local ethical approval (MRC Harwell AWERB committee). At Hospital Clínico San Carlos (19/017-II, 20/11/2020) and La Laguna University (CEIBA 2016-0194 and CEIBA 2022-3186), all animal procedures were approved by the local ethical committees of animal care and use in accordance with the European and Spanish regulations (2010/63/EU and RD 1201/2005).

#### Mouse line generation and genotyping

As reported previously (Fratta et al., 2018) the M323K line was identified after screening DNA archives from large-scale chemical mutagenesis projects (Acevedo-Arozena et al., 2008; GONDO et al., 2010). This mutation was placed on congenic C57BL/6J and DBA/2J backgrounds to eliminate other potential mutations. Subsequently, the two congenic lines were crossed to produce viable F1 C57BL/6J x DBA/2J homozygotes.

At weaning, all mice were genotyped by PCR from an ear biopsy. We used a multiplexed PCR assay (two primer/probe pairs, Supplementary Table 1) for genotyping with data collected at the end of the qPCR process.

### Phenotyping tests

All tests were conducted in both sexes separated, unless otherwise stated.

#### Weights

Non-anaesthetised mice were weighed inside a beaker monthly and before the appropriate behavioural test. To correct for animal movements, an average weight was taken after five seconds.

#### Marble burying test

Nine glass marbles were evenly spaced in rows of three in an IVC containing approximately 4 cm depth of aspen bedding. The mouse was placed in the cage and IVC lids were positioned on top for 30 minutes. For the total score of buried marbles, the following scores were given less than 25% buried 0, around 50% buried 0.5 and more than 75% buried 1. The score was then normalised to the total number of marbles (9). Finally, the score was normalised to the average number of marbles buried by the wild-type at 3 months of age.

#### Nesting test

Mice were placed in individual cages with wood-chip bedding but no environmental enrichment items. Nestlets (3g weight) were placed in the cage overnight. Nests were assessed the next day on a rating scale of 1-5. 1-nestlet not noticeably touched (90% intact). 2-nestlet partially torn (50-90% remaining intact). 3-nestlet mostly shredded but no identifiable nest site. 4-an identifiable but flat nest (more than 90% of the nestle torn). 5-a near perfect nest with walls higher than mouse body height.

#### Novel object recognition test

Initially, the mouse was habituated within the arena (methacrylate box of 40 cm x 40 cm). The habituation process lasted two days and the mouse spent 10 minutes in the box each day. On the third day, the test was carried out by measuring the time spent with each object. To do this, the mouse was placed in the box for 10 minutes with the two initial objects (two equal weights). One of the objects was removed (always the same one) and a novel object (a plastic lemon) was placed in the arena for the 5 minutes of the test. After 15 minutes, the mouse was removed and placed back into its home cage. The Novel Object box was cleaned before each run with 70% alcohol. To quantify the results, the following ratio was used: (time spent with the new object − time spent with the known object)/total time (300 s). Finally, the score was normalised to the ratio of time by the wild-type at 3 months of age.

#### Light-dark box

A high sided Perspex box split into two zones, dark and light, was used in this test. The dark zone had black walls and a black lid. The light zone had clear walls, a 64 white floor and no lid. The dark wall separating the zones had a small opening, allowing unrestricted movement between the zones. Light levels in the testing room (and light zone) were set at 100 lux. At the start of the experiment, the mouse was placed in the light zone and video tracking was initiated. The mouse was removed from the box after 5 minutes. The time spent in each zone and frequency of entries were calculated from the videos using the Ethovision software (Noldus, Netherlands).

#### Fear conditioning test

On day 1, mice were placed individually into square arenas (17×17 cm, height 25cm) with metal grid floors (Ugo Basile, Italy). Light levels were set to ∼26 lux in each arena. A conditioning protocol was initiated using the AnyMaze software (Version 4.5, Stoelting, USA) and consisted of 120s baseline recording, a 20s 90db tone (conditioned stimulus, CS), immediately followed by a 0.5s 0.4mA shock (unconditioned stimulus, UCS). The conditioning test lasted for 4 minutes and 20 seconds. Mice were then removed and placed in the home cage (Nobili et al, 2017). Arena was cleaned with Distel disinfectant and dried between each trial. The conditioning trial was carried out in the morning. On day two, mice were placed back in the same arena to assess response to context. No tones or shocks were administered. This was carried out in the morning. Each trial lasted for 5 minutes. Four hours after the context trial, mice were placed in a new environment to assess response to cue. Round arenas, with plastic floors and stripped walls, were used. The scent of vanillin was rubbed on the sides of the arena. Light levels were reduced to 15 lux. Trial consisted of 120s baseline recording, followed immediately by a 20s 90db tone. The cue trial lasted for 3 minutes. Mice were video tracked using AnyMaze software and freezing time was calculated using AnyMaze freezing detection parameters.

#### Grip strength test

Mice were placed on the grip meter (Bioseb) with all four paws on the grid. Once on the apparatus, traction was applied via the tail. The resistance the mouse applied on the grid was recorded. Each mouse had three runs. The mean value was corrected for the individual animal’s body weight. Finally, it was normalised to the mean strength by the wild-type at 3 months of age.

#### Grid test

For this test, a metal grid at a height of about 30 cm was used. The mouse was held upside down for 1 minute in each of the three repetitions. The first two repetitions served as habituation for the mouse and were performed consecutively. The third was the objective measure of the test and was performed 3 minutes after the end of the second trial. In addition, the mouse was weighed on the day of the test. After the test the animal was returned to the cage with its littermates. Finally, it was normalised to the mean time to fall by the wild-type at 3 months of age.

#### Muscle CMAP (Compound Muscle Action Potential) and Spontaneous Activity Measurements

Pollari’s protocol (Pollari et al., 2018) was followed for the electromyographic studies measurement of the muscle of the mice. An analogue stimulator (Grass Telefactor W. Waarwick, RI USA model 588) and a stimulus isolation unit (SIU) (model PSIU6 (Grass Instrument Co USA)), plus an AC/DC amplifier model 3000 (A-M Systems) were used. The recording interface was CED1401 and the software used for recording and subsequent analysis was Spike v8.04 (Cambridge Electronic Design Limited).

Briefly the mouse was anaesthetised and kept with isoflurane until the procedure was completed. Five electrodes were placed on the mouse: two stimulation electrodes (at the starting and at the end of lumbar region of spinal cord) and two recording electrodes (one in the area near the gastrocnemius and another in the area of the Achilles tendon and the ground electrode on the opposite side of the body). The protocol to register the maximal responses followed increasing intensities of stimulation, starting at 1 mA up to 14 mA. After the procedure, the mouse was left in a warm cage and monitored until it fully recovered, before returning it to its cage. During the following days, the mice were monitored to check health status.

Spike 8.02 software was used to quantify CMAP. To do this, 3 cursors were placed, cursor 0 to mark the event, cursor 1 where the response begins, and cursor 2 where the response ends. Finally, to know the value of CMAP, the highest and lowest value within cursors 1 and 2 was analysed. Finally, it was normalised to the mean CMAP by the wild-type at 3 months of age.

The measurement of spontaneous activity was performed before the CMAP recording. The electrodes were placed on the animal as explained above, we waited 3 minutes without any type of stimulus and recorded the spontaneous response of the mouse. In the case of spontaneous activity, this was analysed by adding two horizontal cursors 1 and 2. The cursors were placed over 20% of the background noise value. The number of spontaneous activity events corresponds to the number of waves that pass over these two cursors, obtaining the total number of spontaneous activity in the 3 minutes of recording.

### Femur Length

Mice were anaesthetised and kept with isoflurane until the procedure was completed. Computed tomography (CT) images were taken in a small animal positron emission tomography (PET)/CT hybrid scanner (Albira ARS, Bruker). CT images were reconstructed applying a filtered back projection. Femur length for each limb was measured in Pmod 4.1 (Pmod Technologies, Bruker) using the data inspector tool. For each animal, average femur length was calculated.

### Immunostainings

Mice were terminally anaesthetised with 0.4 mg/kg of fentanest and 40 mg/kg of thiopental and cardiac perfused with the fixative 4% PFA without methanol in PBS 1X, pH 7.4. The dissected organs were post-fixed in the same fixative overnight at 4 °C and kept on PBS 1X with sodium azide 0.1% until use.

For cryosectioning, tissues were cryopreserved by immersion in 30% sucrose. Tissues were dried and embedded in OCT using isopentane and dry ice, and frozen at −20°C until cut with a cryostat (Leica). Transverse sections at 35 µm were collected free-floating through the lumbar spinal cord and brain frontal cortex. Free-floating sections were then blocked with PTB (PBS 1X, Triton 0.02% and Bovine Serum Albumin (BSA)) and then incubated with the primary antibodies in PBS with Triton 0.02% (for the list of antibodies used, see Supplementary Table 2) and the secondary antibodies (Alexa-Fluor, ThermoFisher), and then mounted together with Hoechst stain to detect nuclei. Images were taken using a confocal microscopy Olympus Fluoview FV1000 with a version system 3.1.1.9.

### Imaging analysis

Images were uploaded into Fiji/ImageJ for analysis and intensity quantification, colocalization and cell number counts. For the total intensity protocol, we converted the image into binary then we quantified the number of particles (analyse particles) by selecting the IntDen function of ImageJ/Fiji. Subsequently, the total number of particles was multiplied by the IntDen value, obtaining the total intensity value. This protocol was used to analyse IBA1, GFAP and Sudan Black Staining. The ‘Colocalization Threshold’ macro in the Fiji software was used to calculate the percentage of colocalization. This protocol was used to analyse TDP-43 colocalization versus Dapi. For this, the images to be analysed were selected in red and green colours, the macro was selected, and the default options were used. The TM1 and TM2 values refer to how much of one protein colocalizes with the other and vice versa. Finally, to analyse the total number of particles, we selected the area of interest or the total area and selected "analyse particles" that have refined the data close to the value of a pixel. This protocol was used to analyse Chat, Ctip2, Lower motor neuron and Parvalbumin staining.

### Sudan Black

Fixed brain cryosections at 35 µm were mounted on slides treated with “TESPA”. Subsequently, they were left to dry for 1 hour at 37°C in an oven. The tissues were immersed in 70% EtOH for 1 minute followed by a completely immersion in Sudan Black 0.5% for 20 minutes. Successive washes were carried out in 70% EtOH until the desired degree of specific staining is achieved. Finally, the slides were washed with distilled water and counter stained with Crystal Violet to label the nuclei for 30 seconds. The slides were then dried and mounted with mounting medium. Images were taken with Grundium Ocus® microscope scanners.

The images were loaded into Fiji/ImageJ for the analysis and intensity quantification. Separated colour layers by ROI followed by the total intensity protocol as before.

### Western blot analysis

Fresh dissected tissues were snap frozen and stored at −80 °C until required. Tissues were homogenised using mechanical disaggregation with beads (Precells) in RIPA buffer (Thermofisher) at 4°C with protease inhibitors cocktail (Roche). After 20 minutes of centrifugation at 13000 g at 4°C, the supernatant was collected, and the total amount of protein was quantified using the DC Assay method (BioRaD). 30 µg of protein was loaded for each of the samples in a 10% acrylamide precast gel (Genescrip, US) under reducing conditions, run and transferred into a PVDF low fluorescence membrane (GE healthcare). After blocking, the membranes were incubated with TDP-43 primary antibodies (see Supplementary table 2), and labelled secondary was added for infrared detection and scanned in the infrared scanner Clx (LiCor). The total protein detection kit was used following manufacturer’s instructions (LiCor) for the loading correction. The intensity of the bands was quantified using the image software Image Studio v.5.2 (LiCor).

### Quantitative PCR analysis

Fresh dissected tissues were snap frozen and stored at −80 °C until required. RNA was extracted from tissues using the RNeasy Lipid Tissue Mini Kit (Qiagen). Using the extracted RNA, cDNA synthesis was performed using the High Capacity cDNA Reverse Transcriptase Kit (ThermoFisher Scientific) with 2µg of total RNA. cDNA for qPCR reactions was used at a final concentration of 20 ng per well. Fast Sybr Green mastermix (ThermoFisher Scientific) was added, followed by the appropriate primer; final well volume of 20 µl. Primers are listed in Supplementary Table 1. All reactions were run in triplicate.

### Structural MRI

Female mice were used for these experiments at 3 months of age (N = 7 *Tardbp^+/+^*, N = 6 *Tardbp^M323K/M323K^*) and at 12 months of age (N = 8 *Tardbp^+/+^*, N = 8 *Tardbp^M323K/M323K^).* Mice were anaesthetised with ketamine/xylazine and intracardially perfused with a first flush of PBS-Gd (Gd: 2mM Gadovist, Gadolinium contrast agent, Bayer Vital GmbH, Germany), and then with 4% PFA-Gd. The brains were obtained with the skull intact and kept for extra 24h in 4% PFA-Gd at 4°C and stored until used in a solution made of PBS with Gd and sodium azide at 4°C. Skulls at 12 months were scanned in a multi-channel 7.0 Tesla MRI scanner (Agilent Inc., Palo Alto, CA) using a custom-built 16-coil solenoid array (36). A T2-weighted 3D fast spin-echo (FSE) sequence with cylindrical k-space acquisition sequence was used (37), with TR = 350ms, TE = 12ms, echo train length (ETL) = 6, effective TE = 30ms, two averages, FOV/matrix-size = 20 × 20 × 25 mm/504 × 504 × 630, and total-imaging-time 14 h. For all samples, the resulting images had isotropic resolution of 40µm. Skulls at 3 months were scanned in a 7-Tesla 306 mm horizontal bore magnet (BioSpec 70/30 USR, Bruker, Ettlingen, Germany) with a ParaVision 6.0.1 console was used to image brains in skulls. Eight samples were imaged in parallel using a custom-built 8-coil solenoid array. To acquire anatomical images, the following scan parameters were used: T2W 3D FSE cylindrical k-space acquisition sequence, TR/TE/ETL = 350 ms/12 ms/6, TEeff = 30ms, four effective averages, FOV/matrix-size = 20.2 × 20.2 × 25.2 mm/504 × 504 × 630, total-imaging-time = 13.2 h. The resulting anatomical images had an isotropic resolution of 40µm voxels. All images were registered together using pydpiper (Friedel et al., 2014; Nieman et al., 2018). Voxel volumes were estimated from the Jacobian determinants and modelled as a function of genotype and cohort. Both absolute volume changes in mm^3^ and relative volume changes, measured as percentages of the total brain volume, were compared. Differences were considered significant for p < 0.05 after false discovery rate (FDR) correction.

### Motor neuron counts

Lower motor neuron counts were performed following a published protocol (Austin et al., 2022). Spinal cords were removed via hydraulic extrusion. A 1 cm section of the lumbar spinal cord bulge was dissected and embedded in OCT and frozen over isopentane on dry ice. Serial transverse sections (20 μm) were cut from regions L1 to L6 of the lumbar spinal cord and collected onto glass slides. Every third section was analysed leaving a gap of 60 μm between sections and ensuring the same motor neuron was not counted twice. Slides were stained for 20 minutes in 74 Gallocyanin (0.3 g gallocyanin, 10 g chrome alum, distilled water up to 100 ml), rinsed with water, dehydrated and mounted using CV Ultra Mounting Media (Leica Biosystems). Sections were scanned using a Nanozoomer slide scanner (Hamamatsu) at 40x zoom and motor neurons counted. Motor neurons with the following criteria were counted: visible nucleolus, dense, a diameter of >15 μm and visible dendritic branching. About 35-40 sections were counted per animal and the level of the spinal cord standardised by morphological assessment, such that equivalent sections from L4-L5 were included for counting.

Upper motor neurons were counted as previously described (Gautam et al., 2015; Özdinler et al., 2011). Briefly, in three matching cerebral cortex sections a comparable 1.5 mm x 1.5 mm area in the 10X objective field corresponding to the motor cortex was determined (Paxinos & Franklin, 2001). CTIP2+ neurons were counted only if both soma and apical dendrite were visible in the same plane. Total number of CTIP2+ neurons were determined in one field/section in a total of 3 sections (N = 3 mice).

### Neuromuscular Junction counts

Once the lumbrical muscles were extracted, they were kept for 10 min in 4% PFA. They were then washed with 1X PBS for 5 minutes; incubated for 30 min in 2% Triton-100 (Tx-100X) (4°C) in 1X PBS; blocked with 5% BSA and 1% Triton-100 for 45 min at room temperature; and subsequently blocked with MouseOnMouse (MOM) in 1X PBS and left for 30 min at room temperature after initial blocking. Once finished, tissues were incubated with primary antibodies in blocking solution. Antibodies: Ms-anti-Synaptophysin at 1:100 and Rb-anti-Neurofilament M at 1:100. The Ac1° were incubated overnight at 4 degrees with slow movement. They were then washed 3 times in 1X PBS for 30 min at room temperature with gentle agitation. Next, incubation was carried out with anti-mouse secondary antibodies, anti-rabbit secondary antibody and 1.5 μg/μl of bungarotoxin labelled in Rhodamine-BGTX in 1X PBS, for 2 hours with gentle shaking and at room temperature. Finally, five 30 minutes washes were performed in 1X PBS. They were mounted with mounting medium (Fluorsave).

### Splicing experiments

Total RNA from Cortex and Spinal cord was isolated using TriZol reagent (Thermo Fisher Scientific) and then retro-transcribed using qScript™ cDNA Synthesis Kit (QuantasBio). PCR for splicing events was conducted using 2X PCR Master Mix (Thermo) with specific primers spanning the differentially expressed exon (Supplementary Table) for 35 cycles. The products were electrophoresed on agarose gels with ethidium bromide (Sigma). Results were analysed using PSI, calculated dividing the intensity of the exon inclusion band by the sum of both exon inclusion and exon exclusion bands. Primer sequence in Supplementary Table 1.

### Statistical analysis

Statistical analysis was conducted using GraphPad Prism and SPSS. Two groups were compared with a single time point using Student’s t-test. Body weights between genotypes, across multiple time-points, were compared using 2-way ANOVA repeated measures/mixed models and Bonferroni correction of multiple testing. Two groups were compared across multiple time points using 2-way ANOVA with Šídák’s multiple comparisons post hoc test. Three or more groups were compared at a single time point using one-way ANOVA with Dunnett’s post hoc test. Statistical analysis of qRT-PCR data was performed on ΔCT values. 2-way ANCOVA (SPSS) was used to correct the energy expenditure data for the lean mass interaction. Statistical significance was defined as p ≤ 0.05 for analysis of phenotyping and molecular biology data. Please see figure legends for sample size - n numbers; n numbers refer to biological samples (i.e. number of animals used in animal experiments). Statistical detail for each experiment can be found in the figure legends. For adipocyte size count statistics, the two-way ANOVA analysis and Tukey’s correction for multiple groups. Where indicated, ∗ = *p* ≤ 0.05, ∗∗ = *p* ≤ 0.01, ∗∗∗ = *p* ≤ 0.001, ∗∗∗∗ = *p* ≤ 0.0001. Statistical analysis and figures of MRI data were performed in RStudio using RMINC, MRIcrotome, data tree, tidyverse, ggplot2, grid, dplyr, viridis, plotrix and graphics packages.

All graphs were generated using GraphPad Prism 9. Venn diagrams were created using Venny2.1. BioRender.com was used to create original diagrams/figures.

## Results

### 1. TDP-43-M323K mutation in homozygosity affects body size and composition

We have shown that adult mice homozygous for the *TDP-43-M323K* mutation are not viable on a pure C57BL/6J background, but are viable on a hybrid C57BL/6J-DBA/2J (B6-DBA) background (Fig. 1A) (Fratta et al., 2018). While hybrid breeding enabled viable homozygous offspring, we report here a slight reduction in the number of homozygous mice produced, deviating from expected Mendelian ratios (29% *Tardbp^+/+^*, 52% *Tardbp^M323K/+^*and 19% *Tardbp^M323K/M323K^,* with a p-value of a Chi Square analysis p<0.0001) (Fig. 1B). Only genetically homogeneous F1 hybrid B6-DBA animals were used in this study, with the expected proportion of males and females produced (with a p=0.2207 in the Chi Square Analysis) (Fig. 1C). We focused on wild-type and homozygous littermates, excluding heterozygous mice as they showed no overall phenotypes in our previous work (Fratta et al., 2018). Both male and female *Tardbp^M323K/M323K^* mice weighed less than their wild-type littermates from birth (Fig. 1D and F), although the difference was only significant for females up to 12 weeks of age. This was accompanied by smaller body size at 9 months of age, as assessed by nose to tail measurements in males (Fig. 1E). As the nose to tail measurement was more variable and inconsistent in females, we performed a more objective valid estimation of body size, measuring the femoral bone length by x-ray. This analysis confirmed that the body size of the *Tardbp^M323K/M323K^* female mice were also smaller than their wild-type littermates (Fig. 1G).

**Figure 1.**
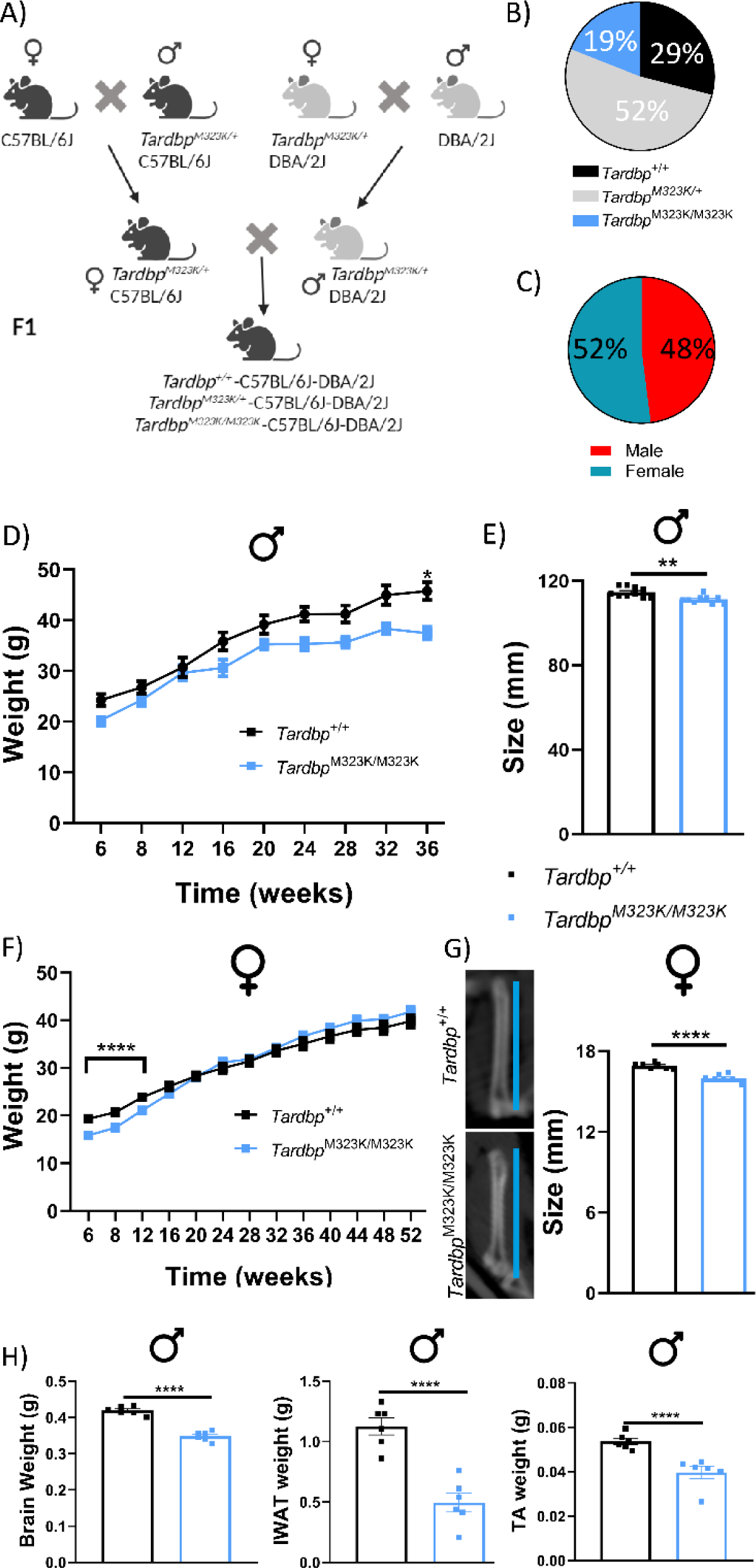
*Tardbp*^M323K/M323K^ mice are developmentally smaller with a sex difference. **A.** Schematic representation of the crosses for the generation of viable homozygous mice. Heterozygous mice from the two congenic lines were crossed to produce viable C57BL/6J-DBA/2J F1 homozygotes. **B and C**. Pie charts showing the genotype (B) and sex (C) ratios obtained in the crossbreeding of mice. **D**. Graph showing monthly body weight measurements from 6-weeks old onwards in male mice. **E.** Measurements of nose-to-tail length in male mice at 9 months of age (*Tardbp*^+/+^ 111.2 mm ± 1.01 and *Tardbp*^M323K/M323K^ 107.6 mm ± 0.38) (for D and E: N =10 *Tardbp*^+/+^; N =9 *Tardbp*^M323K/M323K^). **F.** Graph showing monthly body weight measurements from 6-weeks old onwards in female mice: N = 29 *Tardbp*^+/+^, N = 32 *Tardbp*^M323K/M323K^. **G.** Representative pictures of the femur bone length of *Tardbp*^+/+^ and *Tardbp*^M323K/M323K^ female mice at 1 year old (*Tardbp*^+/+^ 16.92 mm ± 0.09 and *Tardbp*^M323K/M323K^ 15.98 mm ± 0.11), N = 8 *Tardbp*^+/+^, N = 8 *Tardbp*^M323K/M323K^. **H.** Graphs showing smaller the weight of dissected organs: Total brain: *Tardbp*^+/+^ 0.418 g ± 0.005 and *Tardbp*^M323K/M323K^ 0,348 g ± 0.005), muscle (TA, tibialis anterior: *Tardbp*^+/+^ 0.0537 g ± 0.001 and *Tardbp*^M323K/M323K^ 0.3967 g ± 0.002), and fat (iWAT, inguinal White Adipose Tissue: *Tardbp*^+/+^ 1.126 g ± 0.07 and *Tardbp*^M323K/M323K^ 0.4947 g ± 0.08), from male mice at 9 months of age. N=6 *Tardbp*^+/+^, N = 6 *Tardbp*^M323K/M323K^. Data are shown as the mean ± S.E.M. and were analyzed using 2-way ANOVA followed by Tukey multiple comparisons test or Chi-Square test. \**\*p<*0.01, \*\*\**\*p<*0.0001.

We then looked at other organs size and weights, in particular the brain, and found that the *Tardbp^M323K/M323K^* female (data not shown) and male mice had reduced overall brain, fat depots and tibialis anterior muscle weights compared to their control littermates (Fig. 1H). Thus, *Tardbp^M323K/M323K^* mice had reduced body and brain sizes, in both males and females.

### 2. Tardbp^M323K/M323K^ mice display early structural brain changes

The differences in brain weight and size in homozygous mutant mice could be proportional to their smaller body size, affecting all brain areas equivalently whilst maintaining normal structure and function, or could be the consequence of specific neurodevelopmental structural alterations caused by the mutation particularly affecting distinct brain regions. Indeed, previous work in patients (Spinelli et al., 2022) and mouse models have shown that TDP-43 mutations can affect the specific brain regions (Tsuiji et al., 2017). Thus, we performed structural analysis of the brain by magnetic resonance imaging analysis (MRI scans) on wild-type and homozygous female mice at two time points: 3 and 12 months of age, so, we could determine possible early structural changes and/or progressive degeneration during ageing. Significant volumetric differences were observed from 3 months of age, with female *Tardbp^M323K/M323K^* mice having a reduced total brain volume compared to wild-type female littermates (Fig. 2A). The same volumetric differences were observed at 12 months of age, with no significant changes between the 3 and 12 months (Fig. 2A).

**Figure 2.**
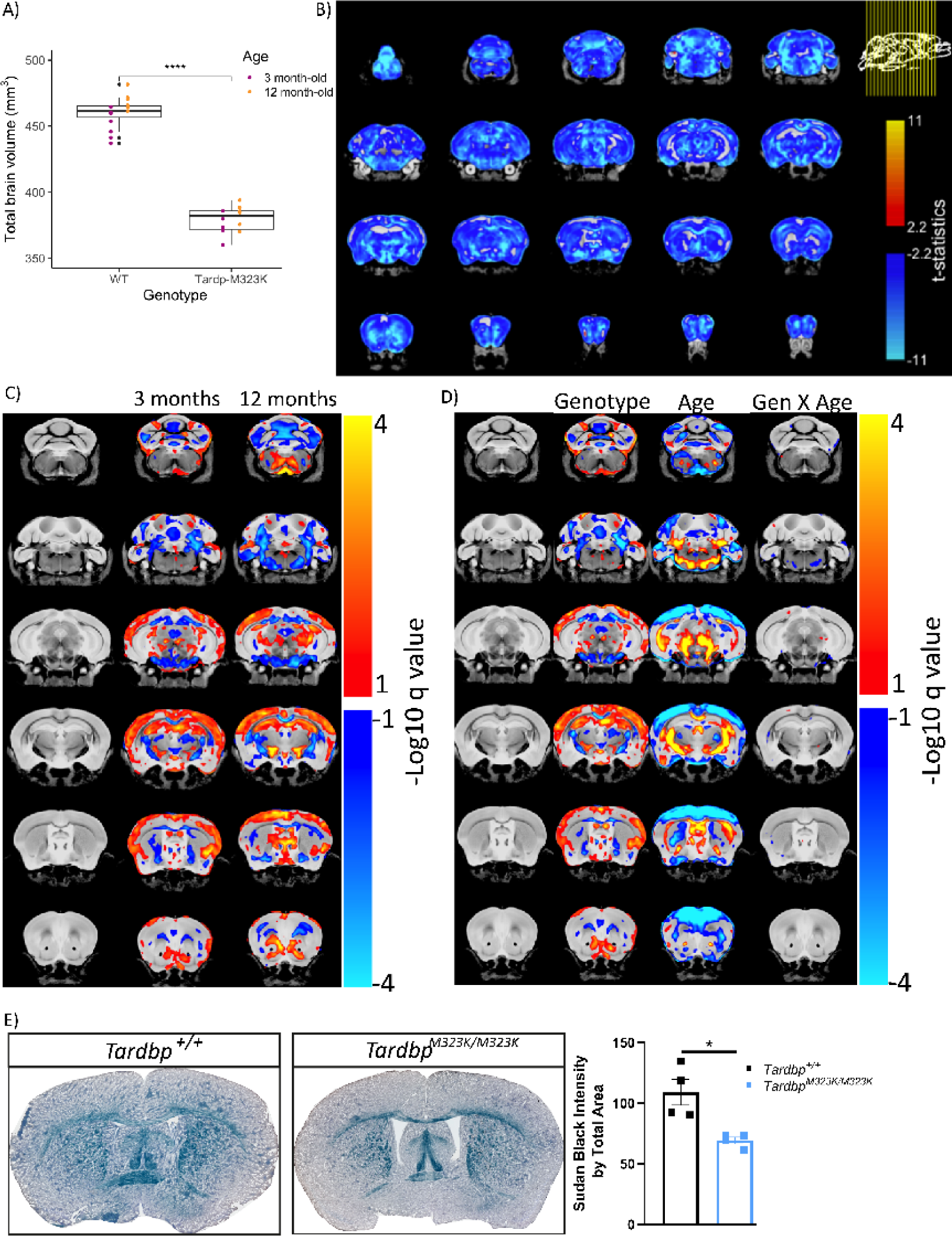
*Tardbp*^M323K/M323K^ mice have relative structural alterations in gray and white matter brain areas from early age. **A.** Graphs showing the volumetric analysis of the brains of female mice at 3 and 12 months of age. At 3 months, N = 7 *Tardbp^+/+^*, N = 6 *Tardbp^M323K/M323K^*, and at 12 months, N = 8 *Tardbp^+/+^*, N = 8 *Tardbp^M323K/M323K^*. **B**. Representative images of the absolute volume of different areas by voxel-based morphometry at false discovery rate (FDR) 5% of the *Tardbp^M323K/M323K^* related to *Tardbp^+/+^*littermates at 12 months of age. **C**. Representative images showing the changes at relative volume between mutant mice at 3 and 12 months of age. **D**. Representative images of the analysis of the separated effect of the genotype and age, as well the combination of the two (genotype X age), at FDR 10%. Blue denotes reduction in voxel size in *Tardbp^M323K/M323K^*mice compared to *Tardbp^+/+^*. Red/Yellow denotes an increase in voxel size. **E.** Representative images of Sudan Black staining of white matter in 3 months-old female mice and the quantification of the whole staining corrected by total brain area. Data are shown as the mean ± S.E.M. and were analyzed using a 2-way T-student test. N = 4 *Tardbp^+/+^*, N = 4 *Tardbp^M323K/M323K^*. *\*p<*0.05, \*\*\**\*p<*0.0001.

In the voxel-wise analysis, almost all brain regions were smaller in *Tardbp^M323K/M323K^* when comparing global volumes calculated from the absolute values of the Jacobian determinants (Fig. 2B and Supplementary Fig. 2A). However, after correcting by total brain volume, the relative volume changes showed that homozygous mutant mice displayed a mixture of relative atrophy and hypertrophy; widespread white matter volume reduction but enlarged hippocampal and cortical regions versus controls (Fig. 2C and Supplementary Fig. 2B). The relative brain volumes at 12 months of age showed similar patterns to those observed at 3 months of age (Fig. 2C and Supplementary Fig. 2C). Only a very small number of voxels presented significant interaction between genotype and age, indicating that the majority of these volumetric changes were present from the early time-point, and were not progressive, pointing to potential neurodevelopmental effects of the mutation (Fig. 2D). Still, there were some areas that showed progressive atrophy including the anterior commissure, which are white matter fibre bundles that connect the brain hemispheres, the amygdala, and temporal lobes. One major finding of the MRI analysis was a general reduction in white matter in homozygous mutants from 3 months of age, we therefore decided to corroborate those results by staining the myelin directly on the brain slices at 3 months of age, finding a clear reduction in the myelin levels in the homozygous mutant brains compared to wild-type (Fig. 2E).

### 3. Tardbp^M323K/M323K^ mice develop cognitive and behavioural deficits from 3-months of age

We performed behavioural tests to investigate the effect of the mutation on cognitive function. Initial observational tests revealed innate behavioural deficits and apathy in *Tardbp^M323K/M323K^* mice in both sexes. From 3 months of age, *Tardbp^M323K/M323K^* mice had lower nesting scores (Fig. 3A). These mice also tended to bury fewer marbles from 3 months of age, and these cognitive deficits worsened at 12 months of age (Fig. 3B and supplementary Fig. 2A). Similarly, early problems in learning and short-term memory were observed in *Tardbp^M323K/M323K^*mice and were also progressively worse with time (Fig. 3C and Supplementary Fig. 2B). Light-dark box and fear conditioning were evaluated in female *Tardbp^M323K/M323K^* mice at 3 and 12 months of age, and indicated anxiety-like phenotype and impaired context learning in mutants only at later timepoints (Fig. 3D, and 3E).

**Figure 3.**
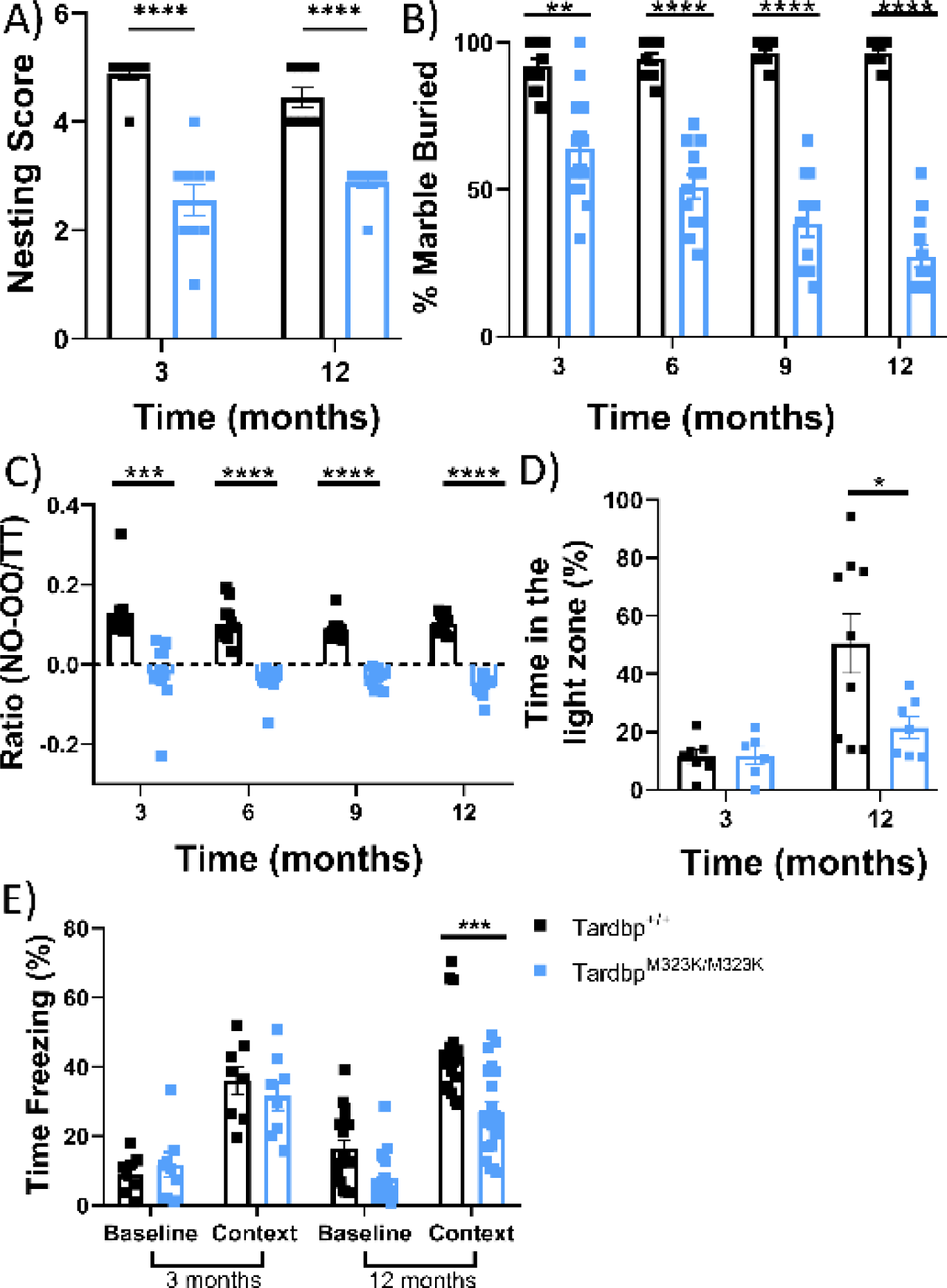
M323K mutation causes early and progressive cognitive and behavioural alterations. All female mice tested longitudinally at different timepoints as depicted in each graph. **A**. Graph showing the capacity of the mice to make a nest. N = 9 *Tardbp^+/+^*, N = 9 *Tardbp^M323K/M323K^*. **B**. Graph showing the percentage of marbles buried. N = 12 *Tardbp^+/+^*, N = 12 *Tardbp^M323K/M323K^*. **C**. Graph showing the ratio of the Novel object recognition (NOR) test. Ratio was calculated as (time on the new object minus time on the known object) divided by the total test time (300 sec) N = 12 *Tardbp^+/+^*, N = 12 *Tardbp^M323K/M323K^*. **D**. Graph showing the percentage of the time spent in the light zone. At 3 months, N = 7 *Tardbp^+/+^*, N = 6 *Tardbp^M323K/M323K^*, and at 12 months, N = 9 *Tardbp^+/+^*, N = 7 *Tardbp^M323K/M323K^*. **E**. Graph showing the percentage of the time the mice are freezing against stimuli at one year of age. At 3 months, N = 8 *Tardbp^+/+^*, N = 8 *Tardbp^M323K/M323K^*, and at 12 months, N = 18 *Tardbp^+/+^*, N = 20 *Tardbp^M323K/M323K^*. Data were analyzed using two-way ANOVA. Data in graphs represent the mean ± S.E.M. *\*p<*0.05, \**\*p<*0.01, \*\**\*p<*0.001, \*\*\**\*p<*0.0001.

Overall, the behavioural analysis revealed that some of the cognitive deficits were present from the earliest time points and were likely mediated by the brain structural alterations identified, whereas other deficits showed a clear progression with age.

### 4. Progressive cellular changes in the brain of Tardbp^M323K/M323K^ mice

We hypothesised that there could be progressive cellular loss and molecular alterations that could underpin the progressive phenotypes, which could not be detected by the MRI scans. First, we evaluated the inflammatory process, which is classically characterised by an increased reactivity of microglia. Thus, we examined the microglia marker IBA1 at 3 and 12 months of age and found a progressive increased in number and intensity of positive IBA1 cells mainly in the white matter areas such as the corpus callosum in *Tardbp^M323K/M323K^* female mice (Fig. 4A). Similarly, we found progressive activation of astroglial cells, using a GFAP marker, in the corpus callosum of the *Tardbp^M323K/M323K^* female mice (Fig. 4B). Interestingly, those areas were also affected in the MRI scans, again pointing towards white matter abnormalities in mutant mice.

**Figure 4.**
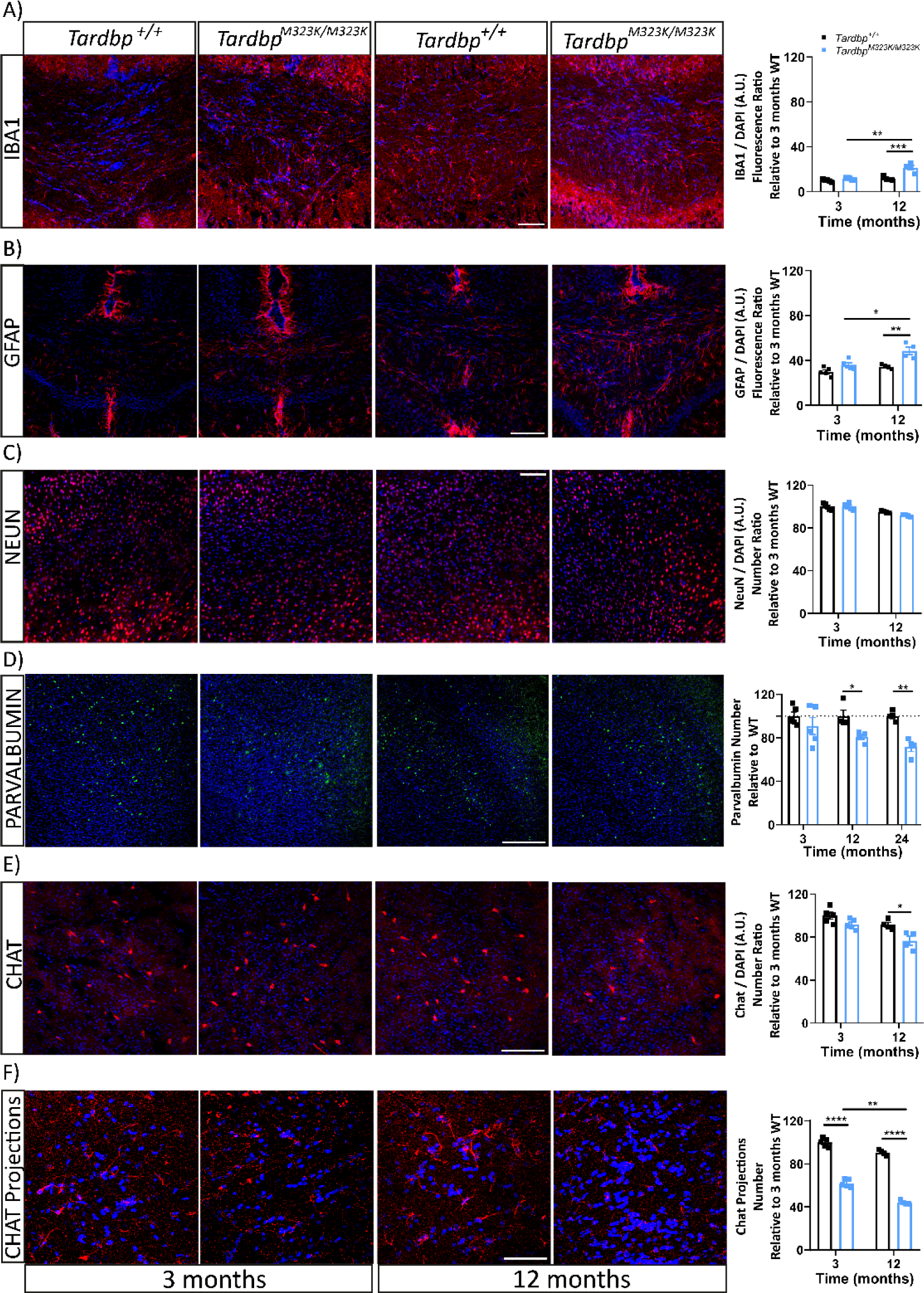
M323K mutation causes progressive cellular losses in different brain areas. Representative Confocal images of the different immunostaining of histological brain sections from female mice at 3 and 12 months of age. All images are merged with the DAPI stained of the nuclei (in blue). **A.** IBA1 staining (in red) at the corpus callosum area. The graph shows the quantification of the total IBA1 intensity in relation to the number of cells (by DAPI) per area, normalized to 3 months *Tardbp^+/+^* (at 12 months of age, *Tardbp*^+/+^ 11.659 ± 0.862 and *Tardbp*^M323K/M323K^ 21.377 ± 2.029) Scale bar: 50 µm. **B.** GFAP staining (in red) at the corpus callosum area. The graph shows the quantification of the total GFAP intensity in relation to the number of cells, normalized to 3 months *Tardbp^+/+^* (at 12 months of age, *Tardbp*^+/+^ 34.434 ± 0.973 and *Tardbp*^M323K/M323K^ 48.594 ± 3.391). Scale bar: 100µm. **C**. nuclear neuronal marker NeuN (in red) in the frontal cortex area. Scale bar: 50 µm. Quantification of the total number of NeuN+ cells per area (data is the average of the three areas analyzed per mouse brain). **D.** Parvalbumin staining (in green) in the frontal cortex region. Quantification of the total Parvalbumin + cells corrected by total number of cells (DAPI) per area, normalized to *Tardbp^+/+^*(until 24 months of age) (at 12 months of age, *Tardbp*^+/+^ 100 ± 5.489 and *Tardbp*^M323K/M323K^ 80.822 ± 2.508, at 24 months of age, *Tardbp*^+/+^ 100 ± 2.268 and *Tardbp*^M323K/M323K^ 71.649 ± 4.192). Scale bar: 100 µm. **E**. Chat staining (in red) at the caudate-putamen region. Quantification of the total Chat+ cells corrected by total number of cells (DAPI) per area, normalized to 3 months *Tardbp^+/+^* (at 3 months of age, *Tardbp*^+/+^ 100 ± 1.797 and *Tardbp*^M323K/M323K^ 62.024 ± 1.720, at 12 months of age, *Tardbp*^+/+^ 90.539 ± 1.366 and *Tardbp*^M323K/M323K^ 44.186 ± 0.933). Scale bar: 100 µm. **F.** Chat staining (in red) at the corpus callosum and primary motor cortex region. Quantification of the total Chat projections and synapsis normalized to 3 months *Tardbp^+/+^* (at 3 months of age, *Tardbp*^+/+^ 100 ± 2.143 and *Tardbp*^M323K/M323K^ 61.024 ± 3.831, at 12 months of age, *Tardbp*^+/+^ 92.525 ± 1.598 and *Tardbp*^M323K/M323K^ 41.186 ± 1.572). Scale bar: 50 µm. Experimental n per time: 3 months N=5 and at 12 months N= 4 per group. Data were analyzed using two-way ANOVA. *\*p<*0.05, \**\*p<*0.01, \*\**\*p<*0.001, \*\*\**\*p<*0.0001.

Next, we checked total neuron numbers with the nuclear NeuN marker in the frontal cortex area and found no differences in wild-type versus homozygous mutant females at 3 or 12 months of age (Fig. 4C), suggesting that the brain structural abnormalities observed did not affect general neuronal numbers, at least in the frontal cortex.

We then decided to assess specific neuronal populations, which could show subtle changes not detected by the general NeuN staining. We measured the number of parvalbumin interneurons in the frontal cortex area of *Tardbp^M323K/M323K^* and *Tardbp^+/+^* female mice at 3, 12 and 24 months of age (Fig. 4D and supplementary Fig. 3A). We observed a progressive loss of parvalbumin interneurons, only evident from 12 months of age. We also looked at cholinergic neurons in the caudate putamen nuclei, another type of interneuron directly related to the motor system, and their projections to the primary motor cortex. The Chat+ stained neurons (soma) in the caudate putamen brain area were progressively decreased between the 3 and 12 months of age in the *Tardbp^M323K/M323K^*mice compared to their wild-type littermates (Fig. 4E). We evaluated the projections of those striatal neurons to the primary motor cortex and found a progressive reduction, which is evident from 3 months of age, even before the loss of the corresponding soma in the striatum (Fig. 4F).

Altogether, these data suggested that the M323K mutation differentially affects the maintenance of specific neuronal populations. The progressive degeneration of these vulnerable cellular populations could be a contributing factor in the progression of some of the cognitive phenotypic alterations identified.

### 5. Tardbp^M323K/M323K^ mice developed progressive motor phenotypes

Our previous characterisation revealed late neuromuscular and motor phenotypes in *Tardbp^M323K/M323K^* mice, with a grip strength deficit observed at 24 months of age in females that was not present at 7 months of age (Fratta et al., 2018). Since we observed clear brain alterations from early ages, we decided to perform a deeper and longitudinal motor characterisation of these mice to determine when motor phenotypes appeared. Motor tests were conducted in both sexes at 3, 6, 9 and 12 months of age. Female *Tardbp^M323K/M323K^*mice developed progressive grid and grip strength deficits from 9 months of age (Fig. 5A, 5B). Interestingly, these deficits were evident earlier in male *Tardbp^M323K/M323K^*mice, at 6 months of age (Supplementary Fig. 4A, B). To evaluate muscle physiological function and also assess potential nerve damage, we carried out electromyographic studies (EMG) in the hind limbs of the same wild-type and homozygous littermates. The amplitude of compound motor action potential (CMAP) can be decreased due to axonal damage in lower motor neurons. CMAP analysis showed a progressive decrease of the muscle response in female *Tardbp^M323K/M323K^* mice from 9 months of age (Fig. 5C), and, again, earlier in males (Supplementary Fig. 4C), suggesting a progressive degenerative process in the motor system in mutant mice. At the same time, we recorded spontaneous muscle activity at 3 and 12 months of age in both sexes, as any involuntary muscle electrical “spontaneous activity” might be pathological (Stålberg et al., 2019). We observed muscle spontaneous muscle activity in *Tardbp^M323K/M323K^* female mice from 3 months, although it was not fully penetrant at this age, with only a few mice showing abnormal activity (Fig. 5D). Interestingly, this phenotype became fully penetrant at 12 months of age, when all the *Tardbp^M323K/M323K^*mice showed spontaneous activity (Fig. 5D), supporting the progressive nature of motor alterations caused by this mutation. Again, these alterations were found earlier in male *Tardbp^M323K/M323K^* mice, as they were already fully penetrant by 9 months of age (Supplementary Fig. 4C).

**Figure 5.**
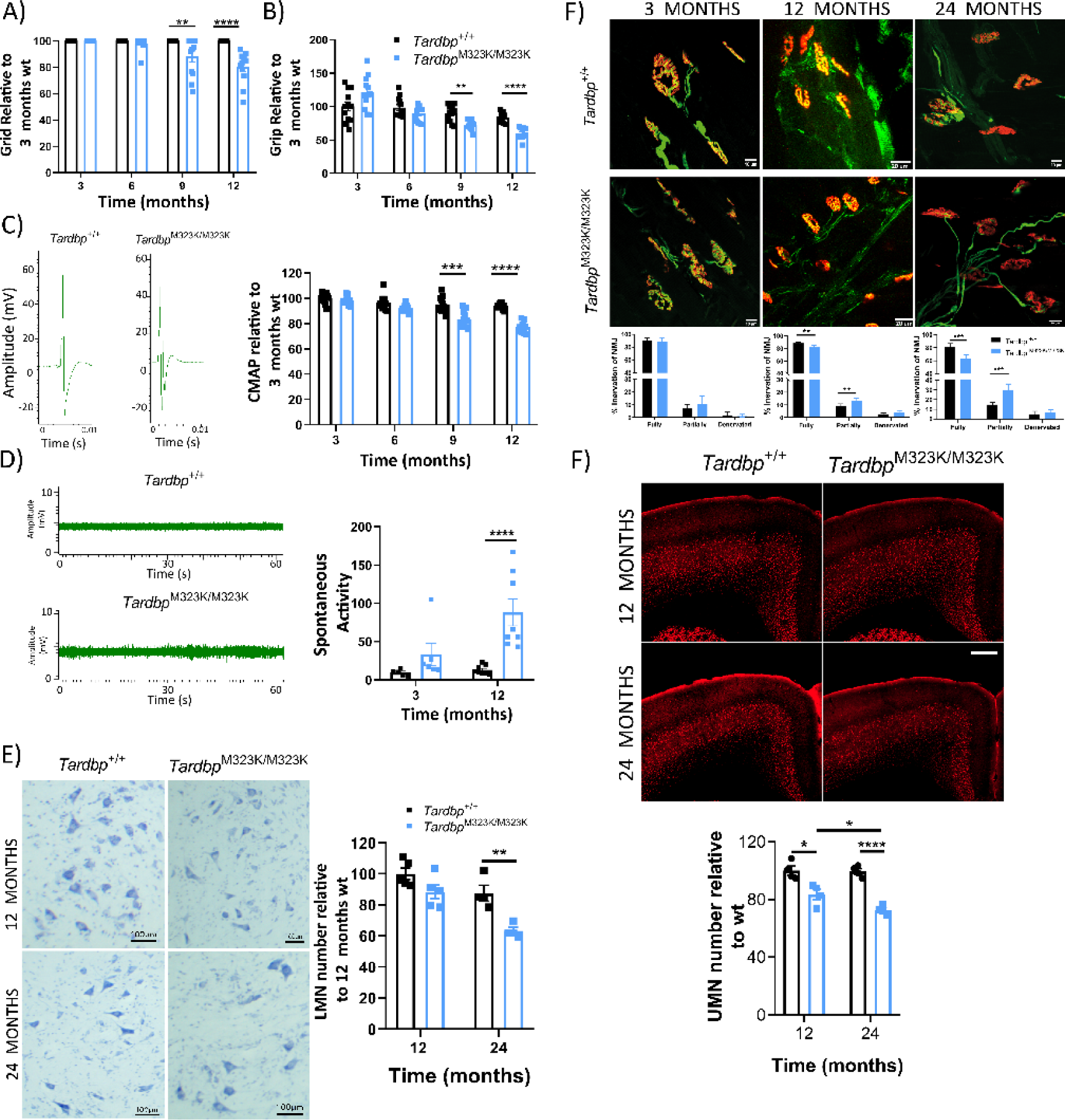
*Tardbp^M323K/M323K^* mice developed progressive motor deficits. Female mice tested longitudinally at 3, 6, 9 and 12 months of age for all motor tests (N= 12 per group). **A.** Graph showing the latency to fall (seconds) in the grid test relative to the 3-month-old *Tardbp^+/+^* data. **B**. Graph showing the corrected means for individual animals’ body weight normalized to 3-month-old *Tardbp^+/+^* data. **C.** Representative images of Compound muscle action potential (CMAP) EMG. Graph showing the CMAP amplitude in the hind limbs relative to the 3-month-old *Tardbp^+/+^* data. **D**. Representative images of Spontaneous Activity EMG. Graph showing the total number of events. **E.** Representative photomicrographs of ChAT^+^ neurons in lumbar spinal cord sections (L1-L5) at 12 and 24 months. N= 4-5 mice per group. Scale bar: 100 µm. Quantification of total ChAT^+^ neurons per area and relative to the 12-month-old *Tardbp^+/+^*data. **F.** Representative confocal images of upper motor neurons (UMN) staining (in red CTIP2) on the primary motor cortex, at 3, 12 and 24 months. N= 4-5 mice per group. Scale bar 100 µm. Quantification of total UMN cells relative to its own wild-type littermates. **G.** Representative confocal pictures of Neuromuscular junction (NMJ) at 3, 12 and 24 months of age. N = 5-6 per gorup. Data were analyzed using two-way ANOVA. Data in graphs represent the mean ± S.E.M. \**\*p<*0.01, \*\**\*p<*0.001, \*\*\**\*p<*0.0001.

Next, we examined whether these early motor deficits were correlated with motor neuron loss in the lumbar region of the spinal cord. In our previous study, motor neuron loss was found at 24 months of age, and we were able to replicate the same findings (Fig. 5E). Here, we also looked at an earlier time point, at 12 months of age, and found no differences in the number of motor neurons between genotypes (Fig. 5E). This result suggests the loss of the lower motor neuron somas might be a later event in the already dysfunctional motor system. Thus, we looked for alterations in the connectivity of these motor neurons with the innervating muscles in hind-limb lumbrical muscles. In agreement with the EMG data, we found a progressive deficit in neuromuscular junctions (NMJ) innervation in *Tardbp^M323K/M323K^*female mice when compared to their wild-type littermates, with a significant increase in partially innervated by 12 months of age, which worsened by 24 months of age, but was not present at 3 months of age (Fig. 5G).

Finally, we examined whether those progressive motor alterations might be initiated in the primary motor cortex in parallel with the observed interneuron neurodegeneration (Fig. 4D-F). Thus, we quantified the number of upper motor neurons (UMN) in *Tardbp^+/+^* and *Tardbp^M323K/M323K^*mice at 12 and 24 months of age by immunohistochemistry of the UMN maker CTIP2 (Arlotta et al., 2005). Interestingly, we found a significant deficit in UMN in *Tardbp^M323K/M323K^* already present at 1 year of age, which progressively worsened by 2 years of age (Fig. 5F).

Altogether, these results show that homozygous M323K mice show truly progressive motor abnormalities that seem to affect the motor cortex and the distal muscles earlier than the lower motor neuron somas.

### 6. TDP-43 functional splicing alterations

From our previous study, we found a gain of splicing function due to the M323K mutation when expressed at endogenous levels, leading to an increase in TDP-43 splicing function in cassette exons known to be modulated by TDP-43, as well as the appearance of pathological splicing events, defined as *skiptic* exons, leading to the exclusion of normally constitutively included exons due to an enhanced TDP-43 repression function (Fratta et al., 2018). Here, we evaluated whether different splicing alterations in skiptics as well as cassette exons known to be modulated by TDP-43 were present in cortex and spinal cord of M323K homozygous mice at different ages. Percentage of exon inclusion (PSI) assessments in a panel of gene exons corroborated previous findings and further showed that splicing function of mutant TDP-43 protein was enhanced in all regions and ages (Fig. 6A-D).

**Figure 6.**
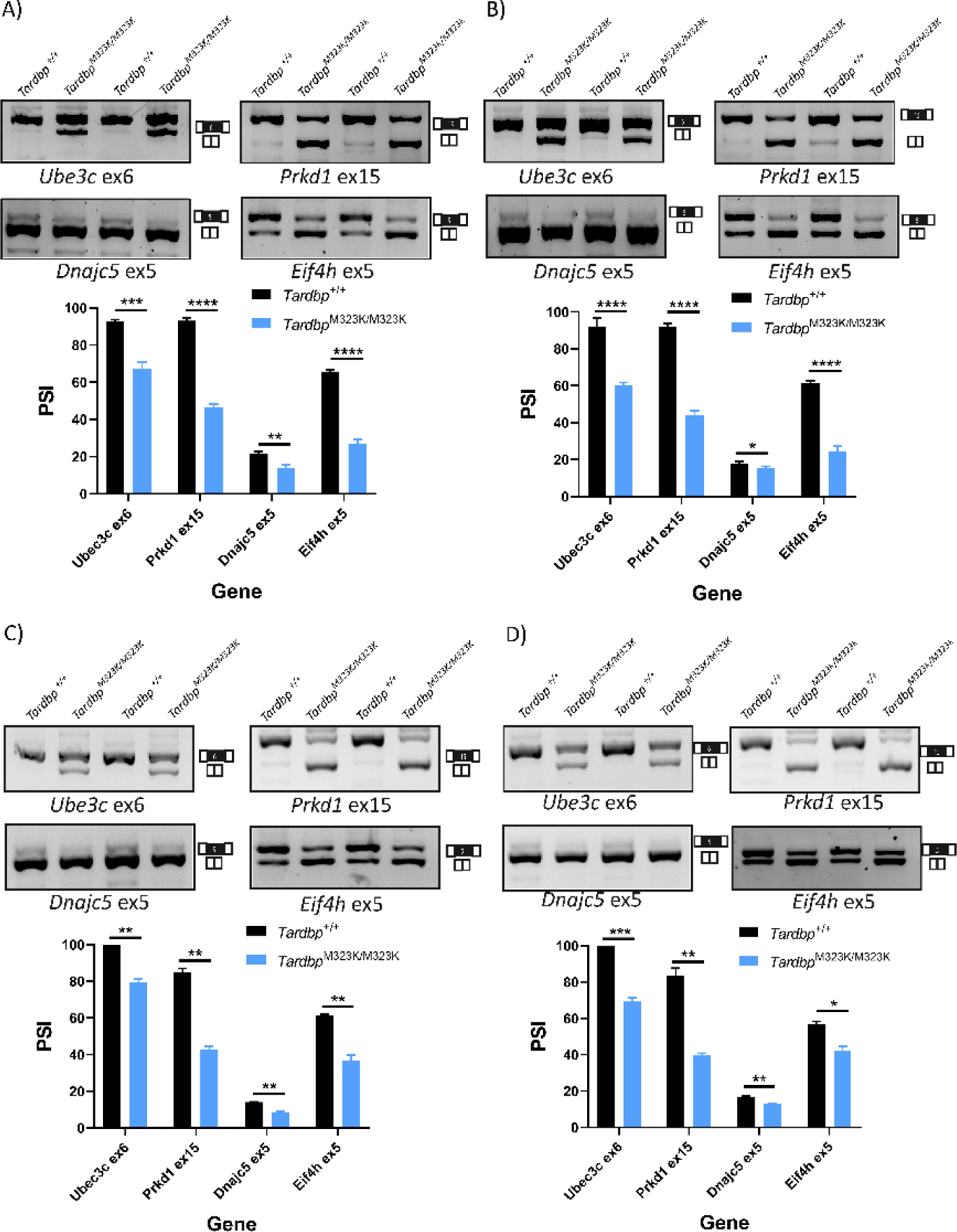
M323K mutation leads to a gain of splicing function in brain and spinal cord. Each panel includes representative images of RT-PCRs run on agarose gels using primers flanking different exons known to be modulated by TDP-43, including two skiptic exons (*Ube3c and Prkd1*) and two cassette exons (*Dnajc5 and Eif4h*). The bands are analysed by densitometry and represented as the percentage of inclusion (PSI) for each exon, depicted as the percentage of the upper band (including the exon) relative to the lower one (exon excluded). **A**. Representative images of RT-PCR and PSI quantification from frontal cortex of 3-month-old M323K mice and littermate controls (N= 4-5 per genotype). **B**. Frontal cortex from 18-month-old (N= 4-5 per genotype). **C**. Representative images of RT-PCR and PSI quantification from the spinal cord of 3-month-old M323K mice and littermate controls (N= 4-5 per genotype). **D**. Spinal cord from 18-month-old (N= 4-5 per genotype). For all panels, mean and S.E.M. are plotted and T-test was used for comparisons, with depicted p-values: *P<0.05; **P<0.01; ***P<0.001; ****P<0.0001.

### 7. M323K mutation led to an increase in TDP-43 protein and mRNA expression accompanied by an increased cytoplasmic mislocalization

Finally, we studied the impact of the TDP-43-M323K point mutation on the expression levels and location of the protein and mRNA *in vivo*, which might explain both the developmental and the progressive alterations observed in homozygous mice.

The M323K mutation is located within the low complexity domain of the TDP-43 protein, known to be important for protein-protein interactions, such as those with other hnRNPs that control TDP-43 splicing function (Buratti et al., 2005) or with chaperones in the cytosol (Lu et al., 2022). Concretely, M323 is located at the N-terminus of the first α-helix of the prion-like region, and the M323 to K mutation changes the polarity of the helix (Fig. 7A). Indeed, we and others have shown that the M323K mutation influences TDP-43 aggregation, showing a higher propensity to form fibrils than wild-type TDP-43, at least *in vitro* (Carrasco et al., 2023; Fratta et al., 2018).

**Figure 7.**
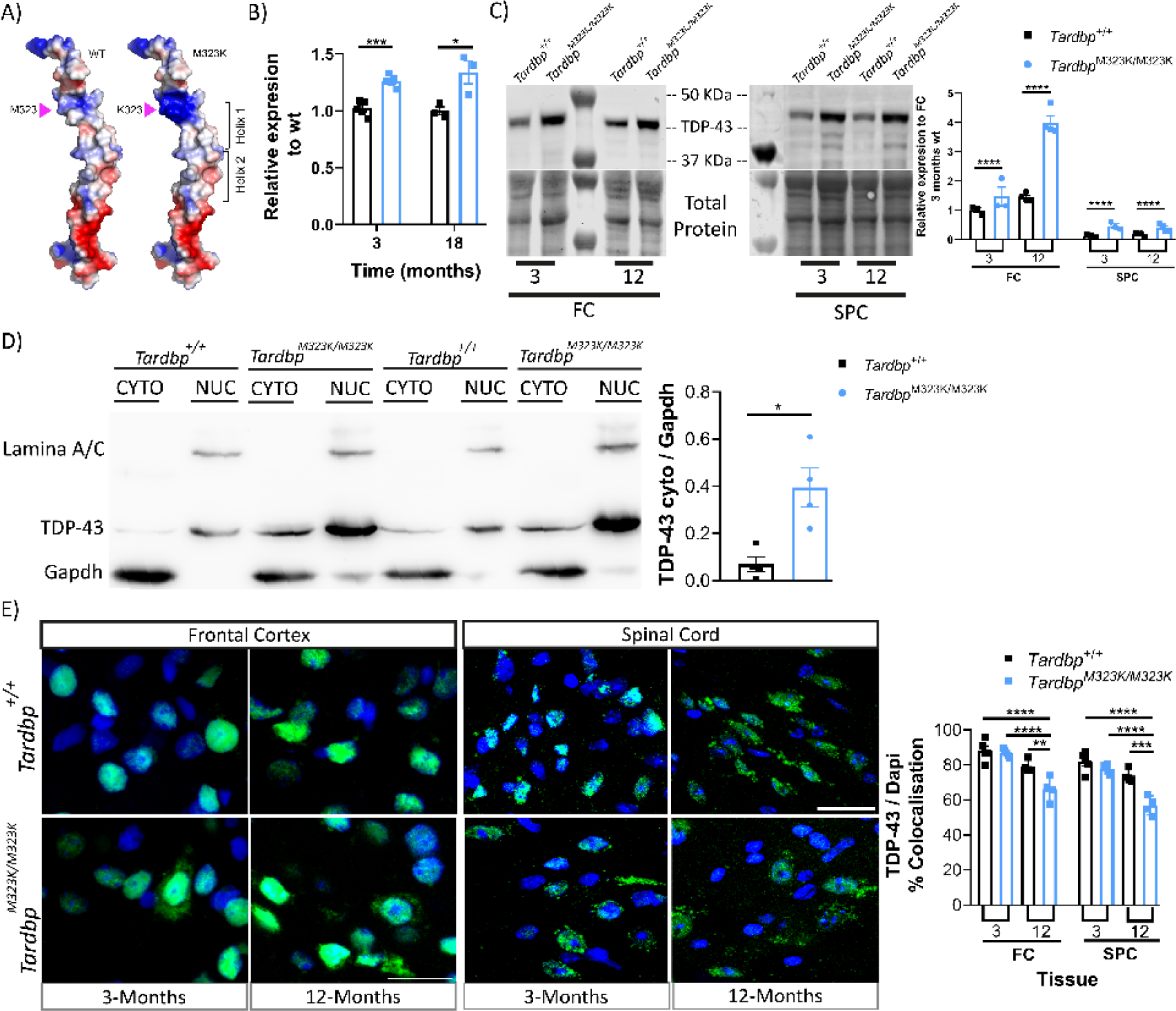
TDP-43-M323K mutant protein had increased expression levels and progressive increased in the cytoplasm location. **A.** Ribbon and electrostatic surface representation of the structure of the wild-type TDP-43 prion-like domain structure (residues 331-360) (Jiang et al., 2016) (left). M323 is shown in sticks mode. M323K point mutation has been introduced to the WT structure to generate the mutant structural model (right). **B**. Gene expression quantification of the *Tardbp* gene in the frontal cortex of female mice at 3 months (N = 5 per group) and 18 months (N = 3 per group). **C**. Representative western blot images of TDP-43 and total protein detection and the quantification relative to the 3-month-old *Tardbp^+/+^* data of the frontal cortex and spinal cord from 3 (N = 3 per group) and 12 (N = 4 per group) month-old female mice. **D.** Representative image of the cytoplasmic/nucleus separation and quantification of the TDP-43 protein at the cytoplasm, normalized to the amount of cytoplasmic GAPDH protein. Frontal cortex of female mice at 24 months of age (N = 4 per group). **E.** Representative confocal images of the TDP-43 immunostaining (in green) merged with nuclear staining DAPI (blue), in the frontal cortex and spinal cord of 3 (N = 4 per group) and 12 (N = 5 per group) month-old female mice. Graph showing the proportion of TDP-43 that colocalizes with DAPI, relative to the FC 3-month-old *Tardbp^+/+^* data. Data were analyzed using two-way ANOVA. Data represent the mean ± S.E.M. *\*p<*0.05, \**\*p<*0.01, \*\**\*p<*0.001, \*\*\**\*p<*0.0001.

We have previously shown that the M323K mutation leads to increased *Tardbp* expression, at least at the transcript level (Fratta et al., 2018). We confirmed this finding in this new cohort at different timepoints, measuring *Tardbp* gene expression levels by qPCR, observing an increased mRNA expression in mutant mice when compared to wild-type littermates in the frontal cortex from 3 months of age, with levels progressively increasing with age (Fig. 7B).

We next looked at the total protein level in different tissues: frontal cortex, spinal cord, skeletal muscle and liver. The total protein levels of TDP-43 were increased in the CNS (frontal cortex and spinal cord) of *Tardbp^M323K/M323K^*mice when compared to littermate controls, with a progressive increase associated with older age in the brain (Fig. 7C, Supplementary Figure A, B). These changes were not observed in peripheral tissues (Supplementary Fig. 6C, D).

To assess if TDP-43 subcellular location was affected by the mutation, we performed subcellular fractionation to separate the nucleus and cytoplasm from brain and spinal cord of wild-type and homozygous mice and found more TDP-43 protein in the cytoplasmic fraction in mutants when compared to wild-type cells (Fig. 7D), an increase that was also progressive with age. To corroborate these findings, we performed immunostaining of the brain and spinal cord of the wild-type and homozygous mice using different TDP-43 antibodies against the N-terminal and C-terminal domains (only showing C-terminal results) in different areas of the CNS. We found similar mislocalization of the TDP-43 protein in the different analysed brain areas, including the frontal cortex and spinal cord (Fig. 7E), as well as, hippocampus and hypothalamus (data not shown). The amount of TDP-43 in the nucleus (measured by the level of colocalization of TDP-43 protein with the nuclear stain DAPI) was reduced by around 20% in *Tardbp*^M323K/M323K^ mice compared to their wild-type littermates at 12 months of age (Fig. 7E), confirming that there was more cytoplasmic TDP-43 in the mutant mice. Interestingly, those changes were not present at 3 months of age, supporting a progressive accumulation of cytoplasmic TDP-43 in M323K mutants.

In summary, these data show a general dysregulation of the TDP-43-M323K protein in the CNS, leading to progressively increased protein levels and cytoplasmic mislocalization that might underlie the progressive phenotypes observed in *Tardbp*^M323K/M323K^ mice.

## Discussion

TDP-43 is of fundamental importance to ALS/FTD, not only because more than 95% of ALS and around 50% of FTD patients present with TDP-43 proteinopathy (Arai et al., 2006; Mackenzie et al., 2007; Neumann et al., 2006), but also because familial ALS and FTD cases can harbour *TARDBP* mutations. The development of sequencing techniques and increasingly accessible screening in the clinic has identified rare *TARDBP* mutations also in clinical neurodegenerative pathologies other than ALS and FTD, such as semantic primary progressive aphasia (svPPA) (Caroppo et al., 2016; Gelpi et al., 2014), flair arms (Moreno et al., 2015), classical Parkinson’s disease and atypical Parkinsonism (progressive supranuclear palsy and corticobasal syndrome) (Chen et al., 2021; Tiloca et al., 2022) and autosomal dominant myopathy (Zibold et al., 2023). Moreover, despite the low prevalence of *TARDBP* mutations, there are a few reported cases of ALS patients carrying homozygous mutations (p.A382T and p.G294V) that do not seem to lead to overtly severe phenotypes (Borghero et al., 2011; Cannas et al., 2013; Corrado et al., 2020; Lombardi et al., 2023). Indeed, patients carrying different *TARDBP* mutations have heterogeneous clinical phenotypes that, at least in the case of ALS, seem to be more often associated with upper motor neuron dysfunction (Lombardi et al., 2023).

Our thorough characterization of M323K homozygous mice has uncovered a myriad of phenotypic alterations including a variety of cognitive as well as motor abnormalities that might model the pyramidal as well as extrapyramidal features seen in the broad clinical outcomes of *TARDBP* mutant carriers. All these imply that TDP-43 also has essential roles outside the motor system, highlighting the importance of studying the wider roles of brain TDP-43 in different model systems.

In this study, we have uncovered a critical role for TDP-43 in brain morphogenesis as evidenced by MRI analysis, showing general and regional brain structural alterations in homozygous mutant mice from an early age. This is in accordance with a previous finding from TDP-43 Q331K KI mice, although the abnormalities found here in M323K mutant mice are more severe (Lin et al., 2021). Overall, this suggests an essential role for TDP-43 in the correct development and maintenance of the central nervous system, as in the case for other hnRNP family proteins, such as FUS (Ali et al., 2023). Interestingly, MRI analysis of mutant *TARDBP* ALS patients have shown that TDP-43 mutations can lead to distinct cortical alterations as well as white matter abnormalities (Spinelli et al., 2022). Indeed, alterations in myelin and oligodendrocytes have been previously reported in other TDP-43 and FUS mouse models (Guzman et al., 2020; Heo et al., 2022; Ho et al., 2021; Wang et al., 2018), and ALS patients (Kang et al., 2013), with premorbid white matter abnormalities recently found in ALS patients (Thompson et al., 2023). Crucially, this is modelled in M323K mice, as this study supports a role for TDP-43 in the development and/or maintenance of white matter, hinting towards alterations in the myelination process that could contribute towards explaining specific cognitive and functional alterations in mutant mice.

Early developmental brain structural abnormalities might underline some of the early deficits that we have found in homozygous animals in phenotypic tests, including nesting behaviour, marble burying, or novel object recognition. However, in other tests such as fear conditioning or light-dark box, deficits do not appear until 12 months of age. For some tests, there is a clear decline with age, for example in numbers of marbles buried, suggesting that at least some cognitive alterations might be underlined by progressive abnormalities that are not developmental. This is indeed also the case at the histological level, as we uncovered a progressive degeneration of parvalbumin and cholinergic interneurons in the cortex, as well as the appearance of astrogliosis after 3 months of age. Some of these progressive changes have also been previously described in transgenic mice overexpressing TDP-43 where a progressive age-associated loss of GABAergic parvalbumin interneurons was reportedly associated with the appearance of TDP-43 inclusions (Tsuiji et al., 2017). Here, in agreement with previous reports on other Tardbp KI strains, we found that interneuron degeneration can occur without TDP-43 inclusions.

This progressiveness is also seen at the motor level, with homozygous mutant mice only showing any deficits after 6 months of age. Critically, these motor abnormalities are underlined by a progressive loss of upper and lower motor neurons, the cardinal pathological hallmarks of ALS. Interestingly, the progressive loss of upper motor neurons appears earlier than that of lower motor neurons, modelling the more pronounced upper motor neuron involvement seen in patients carrying *TARDBP* mutations, as well as in other TDP-43 mouse models (Reale et al., 2023). Thus, different contributors might explain the appearance and progression of motor phenotypes. They could be associated to the progressive loss of upper motor neurons in the primary motor cortex, but also might be related to both, the loss of other parvalbumin GABA interneurons (White et al., 2018) as well as the loss of striatal cholinergic interneurons and their corticostriatal projections to the primary cortex (Maltese et al., 2014; Poppi et al., 2021), indicating that the motor alterations seen in the homozygous mutant mice could be initiated at the primary cortex. Also, distal events initiated at the neuromuscular junction leading to functional muscle alterations at the EMG level are also likely to play a pivotal role. As *Tardbp* Q331K KI mutant mice had reported alterations in the number of parvalbumin interneurons (White et al., 2018) but not underlying lower motor neuron degeneration, possibly other factors might be a play in M323K mutant mice.

TDP-43 levels increase with the normal ageing process in humans (Koike et al., 2021), and although TDP-43 pathology has also been reported in many healthy aged individuals (Uchino et al., 2015), the proportion of people with TDP-43 proteinopathy has been reported to be higher in patients with a neurodegenerative condition. Thus, the progressive nature of some cognitive and all motor phenotypes in M323K mutant mice prompted us to look for underlying abnormalities at TDP-43 expression levels, its function and subcellular distribution in brain and spinal cord at different ages. Indeed, we found an age-related increase in TDP-43 levels in neuronal tissues of aged wild-type mice, and a more pronounced increase in M323K mutant mice. This was accompanied with an increase in TDP-43 cytoplasmic (mis)localization in mutants that was more remarkable in the frontal cortex, which could partly explain some of the progressive behavioural phenotypes in the *Tardbp*^M323K/M323K^ mice. Interestingly, this increase in TDP-43 protein levels and cytoplasmic mislocalization did not correlate with any changes in its splicing function in cortex.

Overall, in this study we take advantage of the M323K mutation, characterizing its wider phenotypes in detail at different levels, identifying different roles for TDP-43 in different regions, and importantly, in different stages of life, emphasizing the need to develop good models to study the complex biology of TDP-43. This is essential to understand TDP-43 pathological mechanisms and their contribution to the heterogeneous clinical alterations in different TDP-43 proteinopathies. The more we understand about the implications of TDP-43 alteration in different clinical disorders throughout the ageing process, the better chances we have to intervene in the different pathological processes with the final aim of providing effective and personalised treatments.

### Inclusion and diversity

We support inclusive, diverse, and equitable conduct of research.

#### Lead contact

Further information and requests for resources and reagents should be directed to and will be fulfilled by the lead contact, Silvia Corrochano and Abraham Acevedo-Arozena.

#### Funding

This research was funded by Consejería de Educación, Juventud y Deporte, Comunidad de Madrid, through the Atracción de Talento program (2018-T1/BMD-10731) and Ministerio de Ciencia e Innovación (PDI2020-1153-70RB-100) to S.C. It was also funded by a Medical Research Council Programme Grant (MC_EX_MR/N501931/1) to E.M.C.F. AA-A is funded by the ISCIII (Miguel Servet fellowship and project grant: PI20/ 00422) and CIBERNED. JMBA is funded by CIBERNED. RMBA is funded by ACIISI. KLM and AMB were supported by the Wellcome Trust (grantsWT202788/Z/16/A, WT215573/Z/19/Z and WT221933/Z/20/Z). The Wellcome Centre for Integrative Neuroimaging is supported by core funding from the Wellcome Trust (203139/Z/16/Z).

## Acknowledgements

We thank the animal care staff in Hospital Clínico San Carlos (Madrid), and the Mary Lyon Centre at the MRC Harwell Institute (Oxfordshire). We thank the genotyping, phenotyping and necropsy core facilities of the Mary Lyon Centre for their help. We are grateful to Jimena Baleriola (Achucarro Basque Center for Neuroscience) for help valuable support with the imaging analysis.

## Declaration of generative AI in scientific writing

The authors declare that there is no AI in scientific writing of this manuscript.

## Credit author statement

**Conceptualization:** TJC, AAA and SC, **Formal Analysis:** JMGC, ZA, JMBA, ABMB, LCFB, PB and PHO **Funding Acquisition:** TJC, AAA and SC **Investigation:** JMGC, ZA, JMBA, IGT, LCFB, JILP, PB, SS, IJC, RMBA, MJSB and RRN **Methodology:** JMGC and JILP **Project Administration:** TJC, AAA and SC **Resources:** TJC, AAA and SC **Supervision:** BJN, JPL, KLM, EMCF, TJC, AAA and SC **Visualization:** JMGC, ZA, JMBA, ABMB and MJSB **Writing-original draft:** JMGC, ZA, JMBA, IGT, EMCF, AAA and SC **Writing-review and editing:** JMGC, EMCF, TJC, AAA and SC

---

**Supplementary Figure 1.**
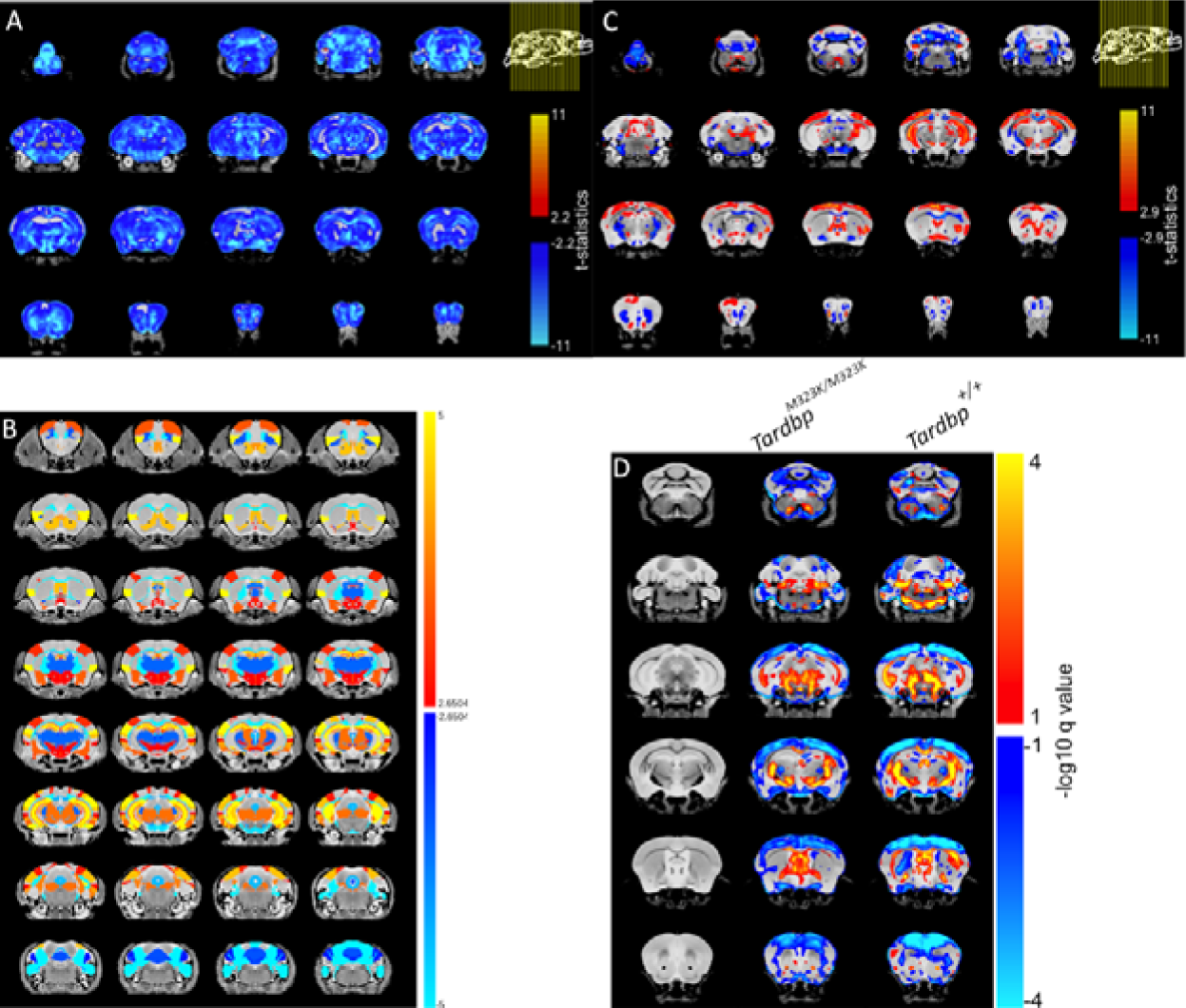
**A.** Decrease in total volume in the whole brain at 12 months of age. **B.** Relative changes in different areas of the brain at 3 months of age. **C.** Relative changes in different areas of the whole brain at 12 months of age. **D.** Changes in different areas of the brain due to the age in wild-type and homozygous mice. (At 3 months, n = 6-7 per group, at 12 months, n = 8 per group).

**Supplementary Figure 2.**
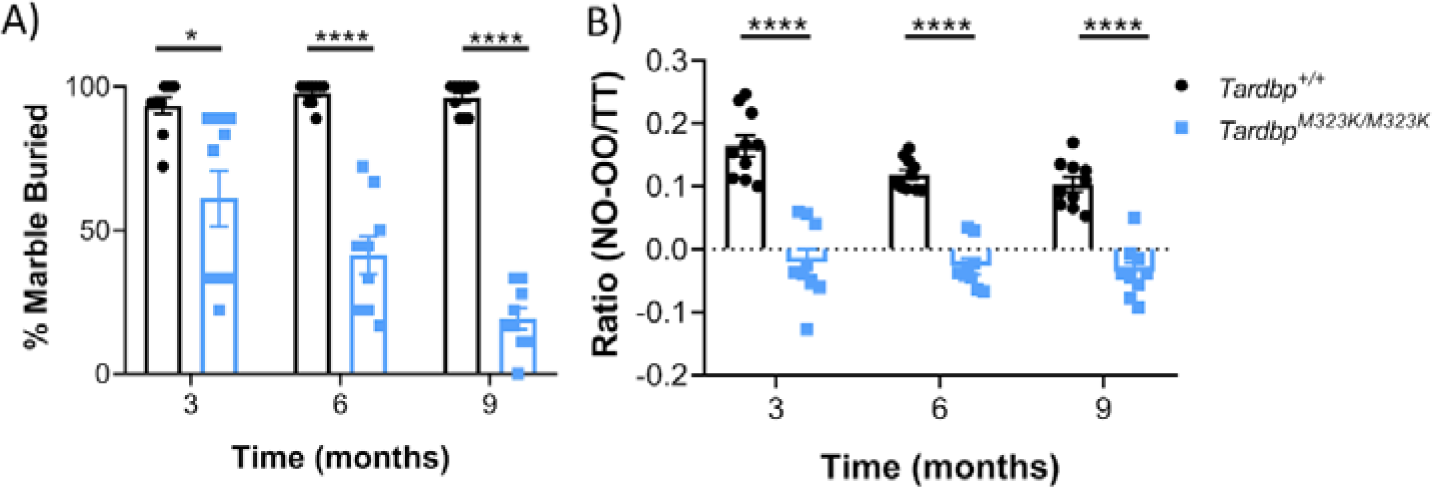
**A.** Marble burying test. The percentage of marbles two-thirds buried was recorded. (n = 9-10 mice per group). **B.** NOR test. To quantify the NOR result, we use a ratio that is calculated as follows: (time on the new object minus time on the known object) divided by the total test time (300 sec) (n = 9-10 mice per group). Data were analysed using two-way ANOVA. Data in graphs represent the mean±S.E.M. *P<0.05, ****P<0.0001.

**Supplementary Figure 3.**
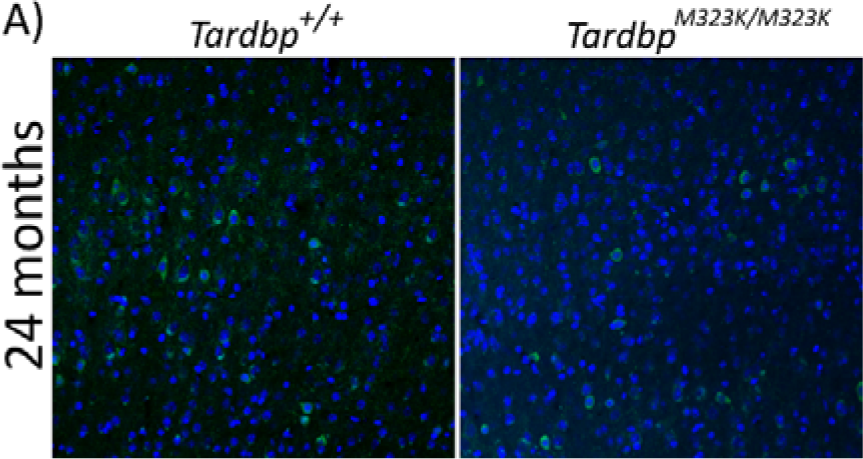
**A.** Representative confocal images of the frontal cortex from female mice at 24 months of age. Merged with Parvalbumin staining (in green) and nuclei (stained with DAPI, in blue).

**Supplementary Figure 4.**
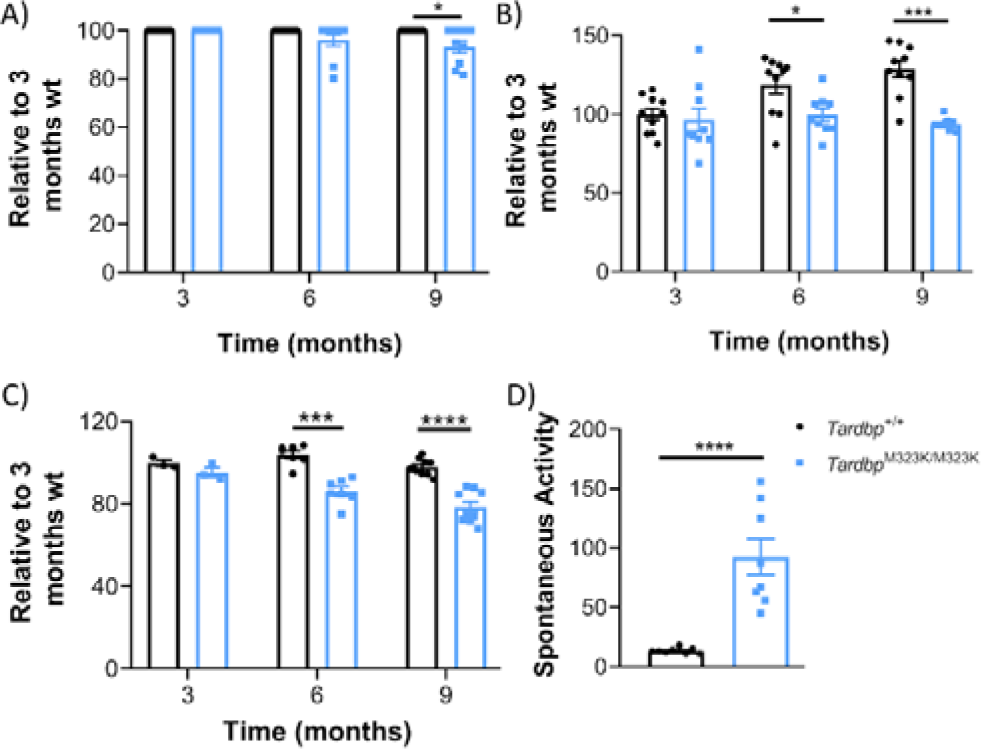
**A.** Grid test. Grid test in *Tardbp^+/+^* and *Tardbp^M323K/M323K^* male mice at 3, 6 and 9 months of age (n = 9-10 per group). **B.** Grip strength test. Corrected means for individual animals’ body weight normalized to 3 months *Tardbp^+/+^* (n = 9-10 per group). **C.** CMAP amplitude in the hind limbs at 3, 6 and 9 months of age. **D.** Spontaneous activity at 9 months of age. Presence of spontaneous activity in *Tardbp^M323K/M323K^* (n = 9-10 per group). *P<0.05, ***P<0.001, ****P<0.0001.

**Supplementary Figure 5.**
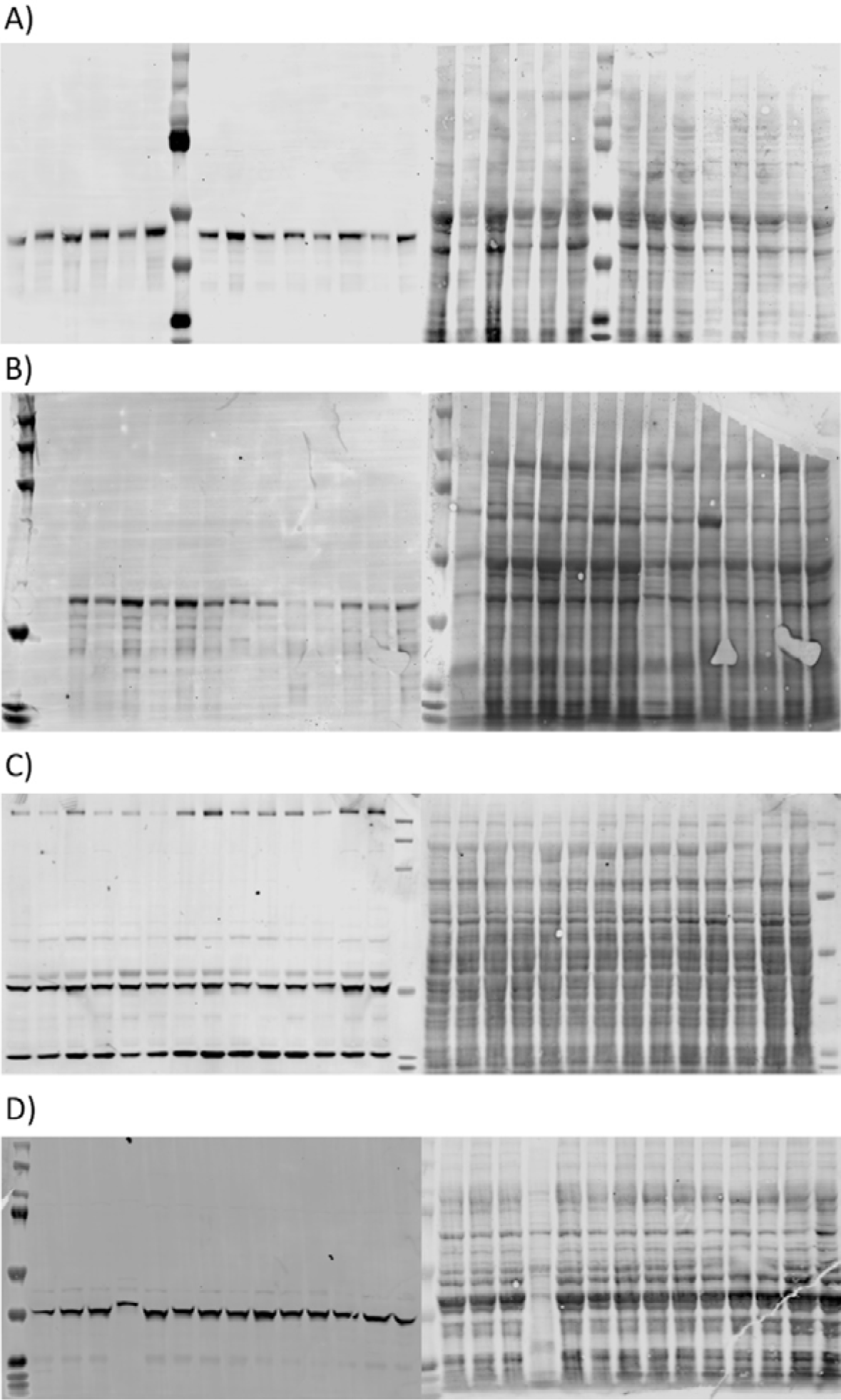
An immunoblot with dilution of 1:1,000 of TDP-43 antibody (ThermoFisher, #MA532627) **A.** Frontal Cortex, **B.** Spinal Cord, **C.** Liver, **D.** Tibialis anterior (n = 3-4 per group and time point).

